# Unmasking Human T Cell Receptor Germline Diversity: 335 Novel Alleles Identified in 47 Pangenome Reference Individuals Using the gAIRR Suite

**DOI:** 10.1101/2025.05.24.655452

**Authors:** Yu-Hsuan Yang, Chi-Yuan Yao, Mao-Jan Lin, Kuan-Ta Huang, Yu-Hui Lin, I-Hsuan Chiu, Sheng-Kai Lai, Chih-Yeh Chen, Ya-Chien Yang, Chia-Lang Hsu, Jacob Shujui Hsu, Chien-Yu Chen, Pei-Lung Chen

## Abstract

The adaptive immune receptor repertoire (AIRR), composed of V(D)J-recombined T cell receptors (TR) and immunoglobulins (IG), is central to adaptive immunity. Accurate AIRR profiling depends on a comprehensive and diverse germline gene set encoding AIRR (gAIRR). Furthermore, gAIRR alleles themselves are associated with immune-related phenotypes and diseases. However, for TR genes, the IMGT database - the primary gAIRR reference - has not been updated since 2020 and lacks sufficient population diversity and complete flanking sequences. To address these limitations, we investigated 47 high-quality genomes from the Human Pangenome Reference Consortium (HPRC) using the gAIRR Suite and identified 335 novel TR alleles - 305 TRV and 30 TRJ - representing 91.6% and 30.9% increases over IMGT records. All novel alleles were crosschecked using two orthogonal pipelines. We also established a comprehensive flanking sequence database, including recombination signal sequences (RSS). We have made all resources publicly available to support immunogenomics research and clinical applications.

## Introduction

The adaptive immune receptor repertoire (AIRR), composed of T cell receptors (TR) and B cell receptors (immunoglobulins, IG), is the cornerstone of the human adaptive immune system. These receptors enable the immune system to mount specific and durable responses to a vast array of antigens, forming the basis for immunological memory and adaptive protection. These receptors are generated through somatic recombination of variable (V), diversity (D), and joining (J) gene segments—a mechanism that produces enormous diversity, enabling immune responses against a wide range of pathogens, tumor cells, and other foreign antigens.^1, 2, 3, 4^ This study distinguishes between the germline gene set encoding AIRR (gAIRR) and its somatically rearranged and expressed counterpart (exprAIRR), which corresponds to the commonly used term “AIRR”. The term exprAIRR is newly introduced in this study to explicitly define the expressed repertoire and avoid ambiguity with gAIRR. While the recombination process eventually links the “core” coding sequences of V(D)J genes, it is regulated by adjacent non-coding flanking elements, particularly recombination signal sequences (RSS). RSS variation can modulate recombination efficiency and thus shape the composition of the immune repertoire.^5, 6, 7^ For instance, an RSS variant enriched among the Navajo population has been linked to heightened susceptibility to *Haemophilus influenzae* infection,^8^ illustrating the functional relevance of genetic diversity in non-coding regions.

Accurate and comprehensive gAIRR references are essential for enabling robust profiling of exprAIRR.^9, 10^ While exprAIRR profiling is widely used to study T and B cell responses in both research and clinical settings, it fundamentally relies on a comprehensive and accurate gAIRR reference.^9, 10^ In addition to serving as a reference for exprAIRR profiling, gAIRR alleles themselves are increasingly recognized as genetic factors associated with disease susceptibility, vaccine response, and adverse immune events. For example, polymorphisms in *IGHV1-69* influence vaccine responsiveness to *influenza* H5N1^11^ and susceptibility to *Staphylococcus aureus*,^12^ *IGHV3-66* variants have been associated with Kawasaki disease.^13^ Similarly, TRBV polymorphisms have been linked to autoimmune diseases^14, 15^ and immune-related adverse events during immunotherapy.^16^ These examples highlight how specific germline variants can directly influence immune phenotypes, making the accurate catalog of gAIRR alleles a critical priority. Despite the importance, our understanding of gAIRR diversity continues to lag behind that of other immune-related gene families such as HLA genes.^17, 18, 19^ At present, the international ImMunoGeneTics information system (IMGT) ^1, 20^ serves as the primary reference database for gAIRR. However, it is increasingly evident that IMGT is incomplete and has not reflected recent advances, especially in the era of massive genomic discovery enabled by next-generation sequencing (NGS). The most recent major update of human TR alleles in IMGT occurred in 2020 (https://www.imgt.org/IMGTgenedbdoc/dataupdates.html), ^1, 20, 21^ highlighting the challenges of keeping pace with NGS-era discovery. More critically, the IMGT database lacks sufficient representation of population diversity, with a clear bias toward individuals of European ancestry.^22, 23, 24^ This underrepresentation limits the global applicability of immunogenomic studies and contributes to disparities in diagnostics and therapeutics.^25^ Compounding this issue is the incomplete nature of flanking sequence data within IMGT. In the current IMGT database, only 65.8% of TRV alleles contain ≥50 base pairs (bp) of upstream and downstream flanking sequences, and some alleles lack this information entirely. These flanking regions—particularly those encompassing RSS—are essential for accurate allele assignment and interpretation of V(D)J recombination dynamics. Their omission can result in ambiguous read alignment and misannotation in gAIRR genotyping and annotation workflows.

Recent advances in NGS and bioinformatics have driven meaningful progress in gAIRR germline databases, especially the IG genes. Long-read targeted sequencing and informatic tools now enable the identification of novel alleles and structural variants.^26^ However, the high cost and technical challenges of targeted long-read sequencing restrict scalability. A promising alternative is the use of publicly available whole-genome long-read sequencing datasets, such as those from the Human Pangenome Reference Consortium (HPRC).^25^ The HPRC has assembled phased diploid genomes for 47 individuals (94 haplotypes) from globally diverse populations using state-of-the-art technologies including PacBio HiFi, Oxford Nanopore, Bionano optical mapping, and Hi-C sequencing. These assemblies are of exceptional quality, with contig N50 values exceeding 80 Mb and structural accuracy surpassing 99%. Importantly, the genomic DNA samples from these individuals are publicly available through the Coriell Institute for Medical Research, making them an invaluable resource for immunogenomic discovery and validation.

To address the current limitations in TR germline allele representation, we applied the gAIRR Suite—a modular bioinformatics pipeline composed of gAIRR-annotate, gAIRR-seq, and gAIRR-call—to the 47 high-quality genomes from the HPRC. This integrative approach enabled the identification of 335 novel TR alleles (305 TRV and 30 TRJ) not previously represented in IMGT (Figure 1a). Notably, we crosschecked the novel alleles through two independent approaches—one based on genome assembly annotation using gAIRR-annotate, and the other using gAIRR-seq for targeted capture-based short-read sequencing followed by allele calling using gAIRR-call. In addition, we extracted and curated the flanking sequences of all the gAIRR alleles from these 47 genomes to create a comprehensive and functionally informative flanking sequence database (Figure 1b). Our work complements recent progress in the immunogenomics field, such as those led by the AIRR Community,^27, 28^ which emphasize rigorous curation and standardized representation of gAIRR reference sets. By contributing high-confidence, population-diverse, and experimentally validated alleles, our findings significantly expand the known TR germline landscape and lay the foundation for future exprAIRR-based research, immunogenetic association mapping, and precision immunogenomics.

**Figure 1.**
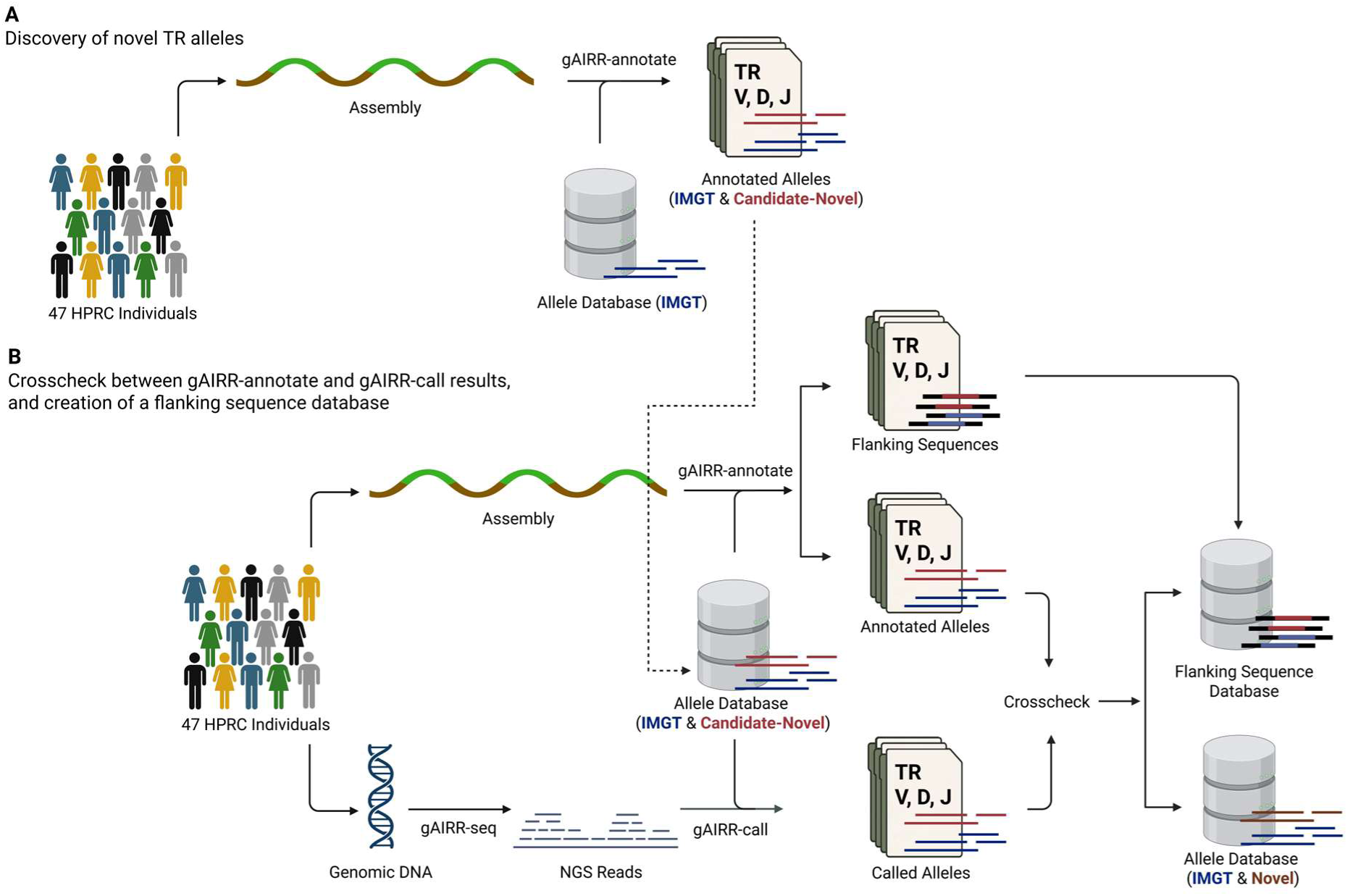
Conceptual workflow of the gAIRR Suite for TR allele discovery, validation, and flanking sequence database construction. (A) Discovery of novel TR alleles. High-quality genome assemblies from the HPRC were analyzed using gAIRR-annotate to identify both reference TR alleles from the IMGT database (blue) and candidate novel alleles (red). (B) Crosscheck between gAIRR-annotate and gAIRR-call results, and creation of a flanking sequence database. This panel illustrates the orthogonal validation process between two independent pipelines. On one side, gAIRR-annotate identifies TR alleles from genome assemblies. On the other, gAIRR-seq captures genomic DNA for targeted sequencing, and gAIRR-call infers alleles from NGS reads. The allele database used by both pipelines includes IMGT reference alleles (blue) and candidate novel alleles (red) discovered in panel (a). Annotated alleles and called alleles are crosschecked to identify validated novel alleles (brown), which are then included in the final allele database. Flanking sequences are extracted to construct a comprehensive flanking sequence database.

## Results

### 1. Identification of 306 novel TR alleles through core sequence analysis

To systematically discover novel TR germline alleles, we applied the gAIRR-annotate tool to the phased diploid genome assemblies of 47 subjects (94 haplotypes) from the HPRC. gAIRR-annotate identifies AIRR alleles by aligning reference sequences from existing germline databases (specifically, the IMGT database) to personal genome assemblies, and it reports novel alleles when mismatches, insertions, or deletions are detected relative to the reference. Using this approach, we identified a total of 309 novel alleles, including 279 TRV alleles and 30 TRJ alleles; no novel alleles were detected in the TRD locus. Each novel allele was assigned a unique identifier using the IgLabel^28^ nomenclature system. For instance, a novel allele identified at the *TRAV1-1* locus with the sequence most similar to the TRAV1-1*01 allele in a specific haplotype was designated as TRAV1-1*01_FYFB, where “FYFB” is the unique identifier assigned by the IgLabel system.^28^

To ensure the reliability of the novel alleles identified using gAIRR-annotate, we performed orthogonal validation by applying the gAIRR-seq/gAIRR-call pipeline to the same 47 HPRC subjects. This pipeline performs targeted capture-based short-read sequencing and independent allele calling. As an illustrative example, subject HG03540 exhibited heterozygosity at the *TRGV3* locus, where both alleles—TRGV3*01_XARG and TRGV3*01_ZZNT—were identified as novel. Sequencing reads aligned exclusively to these two alleles, but not to the corresponding IMGT reference allele, highlighting the power of short-read data to distinguish single-nucleotide differences (Figure 2).

**Figure 2.**
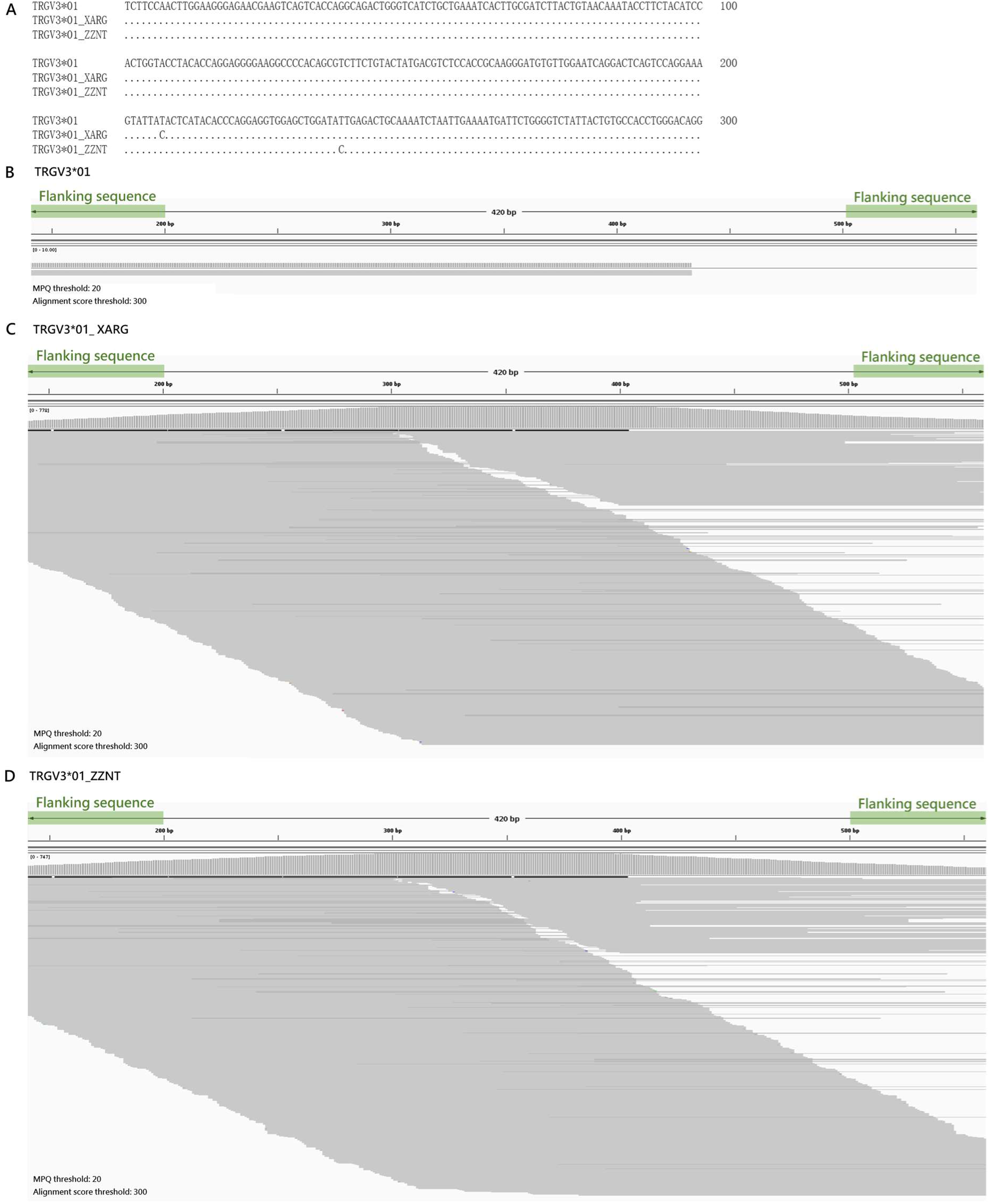
IGV-based validation of novel alleles in HG03540. The *TRGV3* gene in HG03540 is heterozygous, with both alleles being novel (TRGV3*01_XARG & TRGV3*01_ZZNT). (A) Sequence alignment of TRGV3*01, TRGV3*01_XARG, and TRGV3*01_ZZNT. Multiple sequence alignment highlights the variations between reference and variant alleles. (B) IGV visualization using TRGV3*01 as the reference. Only a single read aligns to TRGV3*01, indicating the absence of this allele in HG03540. (C) IGV visualization using TRGV3*01_XARG as the reference. Reads aligned to TRGV3*01_XARG confirm the presence of this novel allele in HG03540. (D) IGV visualization using TRGV3*01_ZZNT as the reference. Reads aligned to TRGV3*01_ZZNT further validate its existence in HG03540. The green-shaded regions indicate flanking sequences, with 60 bp shown on each side of the core allele. Read alignments were filtered using a mapping quality (MPQ) threshold of 20 and an alignment score threshold of 300, ensuring high-confidence, perfect matches to the reference sequences.

Among the 309 novel alleles initially reported by gAIRR-annotate, three alleles— TRBV11-1*01_B6R4, TRGV10*02_FO2S, and TRBJ2-7*01_UZLC—were not confirmed by gAIRR-seq/gAIRR-call. Subsequent investigation attributed these discrepancies to local assembly errors within the genome assemblies (Supplementary Figure 1). After excluding these three cases, a total of 306 novel alleles (comprising 277 TRV alleles and 29 TRJ alleles) were classified as high-confidence.

### 2. Discovery of 29 additional novel alleles through flanking sequence integration

In addition to the core regions of AIRR alleles, flanking sequences play a crucial role in defining precise allele boundaries, underscoring the importance of their comprehensive characterization. The gAIRR-annotate tool is designed to automatically extract 200 bp of flanking sequences on both sides of identified alleles.

Through the integrated analysis of these flanking regions, we identified several cases in which subjects carried alleles that matched known IMGT alleles in their core region but exhibited greater similarity in the flanking regions to other alleles—often those with slightly longer core regions. For example, the IMGT allele TRAV1-1*02 differs from TRAV1-1*01 by a single nucleotide substitution in the middle of the core region (position 199) and a 6 bp deletion at the 3’ end. Initially, gAIRR-annotate classified a novel *TRAV1-1* allele in subject HG002 as TRAV1-1*02, based solely on core-region alignment. However, when additional flanking sequences (50 bp on each side in Figure 3a) were incorporated, alignment analyses showed a more nuanced pattern: the core region aligned with TRAV1-1*02, including the position-199 nucleotide substitution, while the 3′ end—marked by the presence of the 6 bp segment—and the adjacent flanking sequences more closely resembled TRAV1-1*01. This indicates that the HG002 allele should be more accurately classified as a novel variant of TRAV1-1*01 (and we accordingly assigned it as a novel allele, TRAV1-1*01_FYFB), rather than TRAV1-1*02 (Figure 3a). Supporting this interpretation, IGV visualization of gAIRR-seq data (Fig. 3b) demonstrated that sequencing reads aligned perfectly to this novel allele (TRAV1-1*01_FYFB) and not to either of the original IMGT alleles (TRAV1-1*01 or TRAV1-1*02), further confirming its authenticity.

**Figure 3.**
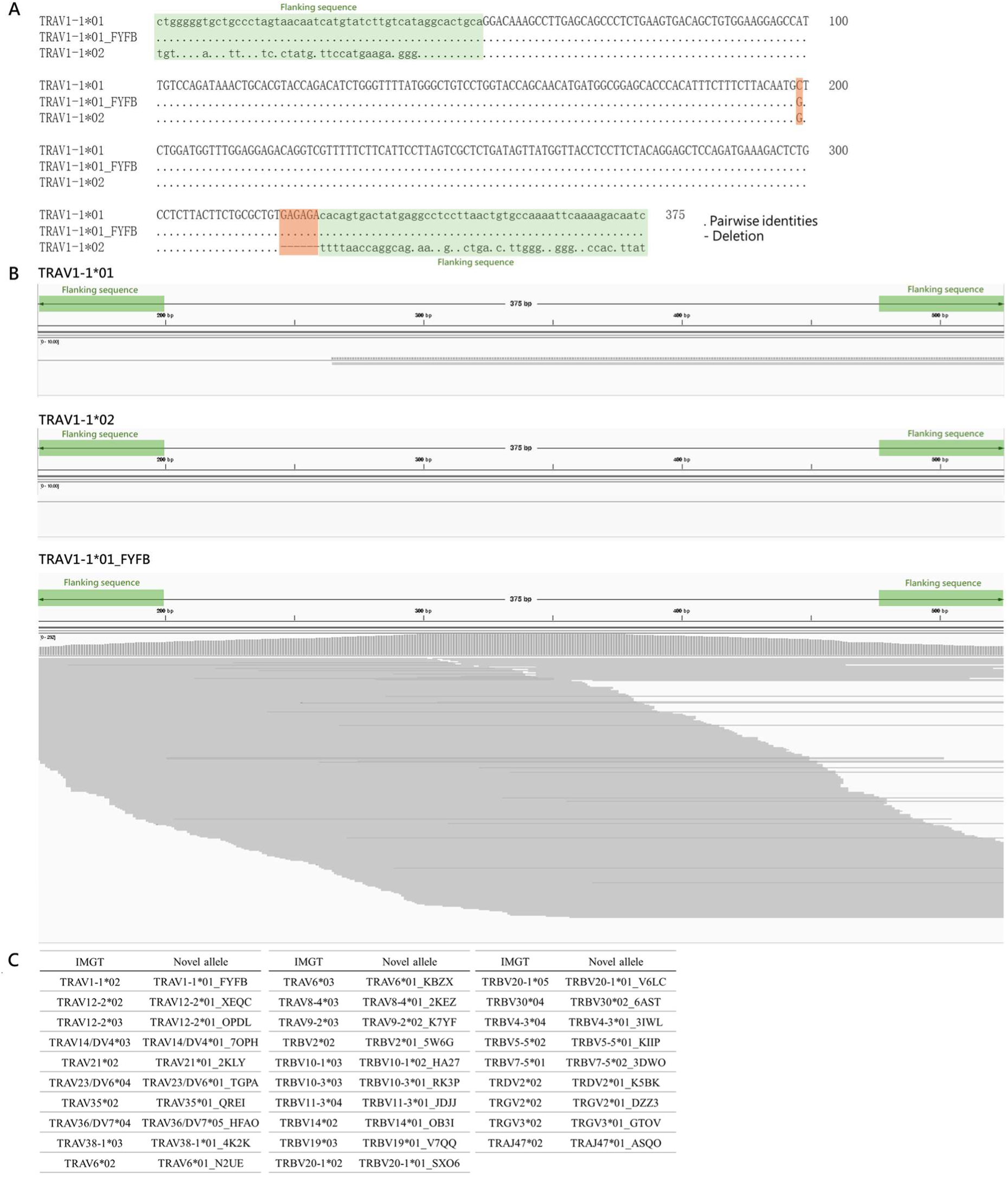
The role of flanking sequences in identifying novel alleles. (A) Impact of flanking sequences on allele classification.TRAV1-1*01 and TRAV1-1*02 differ by a single base change (C to G) at position 199, and TRAV1-1*01 also contains an additional 6 bp sequence (GAGAGA) at the 3’ end of the core region. Initially, gAIRR-annotate identified the *TRAV1-1* allele in subject HG002 as TRAV1-1*02 based solely on the core region sequence (as the G at position 199 in HG002 matches that of TRAV1-1*02). However, when flanking sequences (50 bp on each side, shown in lowercase) were included in the analysis, the *TRAV1-1* allele in HG002 was found to share the GAGAGA sequence with TRAV1-1*01. Therefore, the analysis, including additional flanking sequences, suggests that the *TRAV1-1* allele in HG002 is more similar to TRAV1-1*01 and should be classified as a novel allele of TRAV1-1*01 (TRAV1-1*01_FYFB) rather than a novel allele of TRAV1-1*02. Pairwise identities are indicated by dots (“.”), while deletions are represented by hyphens (“-”). The green-shaded regions indicate flanking sequences, with 50 bp shown on each side of the core allele. Orange boxes highlight the sequence differences between TRAV1-1*01 and TRAV1-1*02. (B) IGV visualization of gAIRR-seq results. Reads from HG002 did not align to TRAV1-1*01 or TRAV1-1*02, despite both covering the core region and flanking sequences. Instead, the reads showed strong alignment to TRAV1-1*01_FYFB. The green-shaded regions indicate flanking sequences, with 60 bp shown on each side of the core allele. Read alignments were filtered using a mapping quality (MPQ) threshold of 20 and an alignment score threshold of 300, ensuring high-confidence, perfect matches to the reference sequences. (C) Novel alleles sharing core sequences with IMGT reference alleles. The 29 novel alleles share core sequences identical to those of IMGT reference alleles, but these sequences are extended and exhibit variations in the flanking regions.

These findings underscore the importance of incorporating flanking sequence information for accurate allele annotation in gAIRR-annotate and any other alignment-based methods. However, an examination of the IMGT database revealed that flanking sequence data remain incomplete: only 65.8% (219/333) of TRV alleles include at least 50 bp of flanking sequences on both sides, and 15 alleles lack flanking sequence data entirely (Supplementary Figure 2a). Consequently, some alleles documented in IMGT might represent incorrect annotations or incomplete sequences, further highlighting the need for thorough validation and potential revision of germline reference databases.

To enhance the sensitivity of gAIRR-annotate for such cases, we implemented a strategy in which an additional 50 bp of flanking sequences on both ends were considered during allele comparison. For IMGT alleles lacking 50 bp of flanking sequence data, we substituted the missing flanking sequences with those from other alleles of the same original gene, under the assumption that alleles of the same original gene are expected to share conserved flanking regions.^29, 30, 31^ After incorporating flanking sequence information into the analysis, we identified 29 novel alleles—28 TRV alleles and 1 TRJ allele—that possessed core regions identical to known IMGT alleles, but exhibited distinct end patterns and flanking sequences matching other alleles (Fig. 3c & Supplementary Figure 2b). All of these 29 novel alleles were further validated using the gAIRR-seq/gAIRR-call pipeline, validating their reliability.

### 3. Crosscheck and concordance across gAIRR-annotate and gAIRR-seq/gAIRR-call pipelines

To ensure the accuracy of novel allele identification, we performed crosscheck between the outputs of gAIRR-annotate (assembly-based) and gAIRR-seq/gAIRR-call (read-based) pipelines. Both tools were independently applied to the same set of 47 HPRC subjects, and allele calls were compared for concordance. A novel allele was considered reliable only if it was supported by both methods. This cross-pipeline validation provides an essential quality control framework for novel allele discovery (Supplementary Figure 3).

Crosscheck between pipelines confirmed 335 out of 338 alleles initially reported by gAIRR-annotate. The three discordant calls, resulting from local assembly artifacts (see Results 1), were excluded from the final dataset.

This high validation rate (99.1%) highlights the robustness of the gAIRR Suite in novel TR allele discovery, as well as the high quality of HPRC personal genome assemblies. The strong concordance between genome assembly-based and targeted sequencing-based approaches supports the reliability of the identified novel alleles and underscores their suitability for incorporation into updated germline reference databases.

### 4. Expansion of TRV and TRJ alleles by 91.6% and 30.9%, respectively, compared to the IMGT database

We identified a total of 335 novel alleles through core region analysis and further refined them via flanking sequence integration, resulting in a substantial expansion of the existing IMGT database. Specifically, the number of TRV alleles increased by 91.6%, and TRJ alleles by 30.9% (Fig. 4a). The full sequences of these novel alleles are provided in Supplementary Table 1.

**Figure 4.**
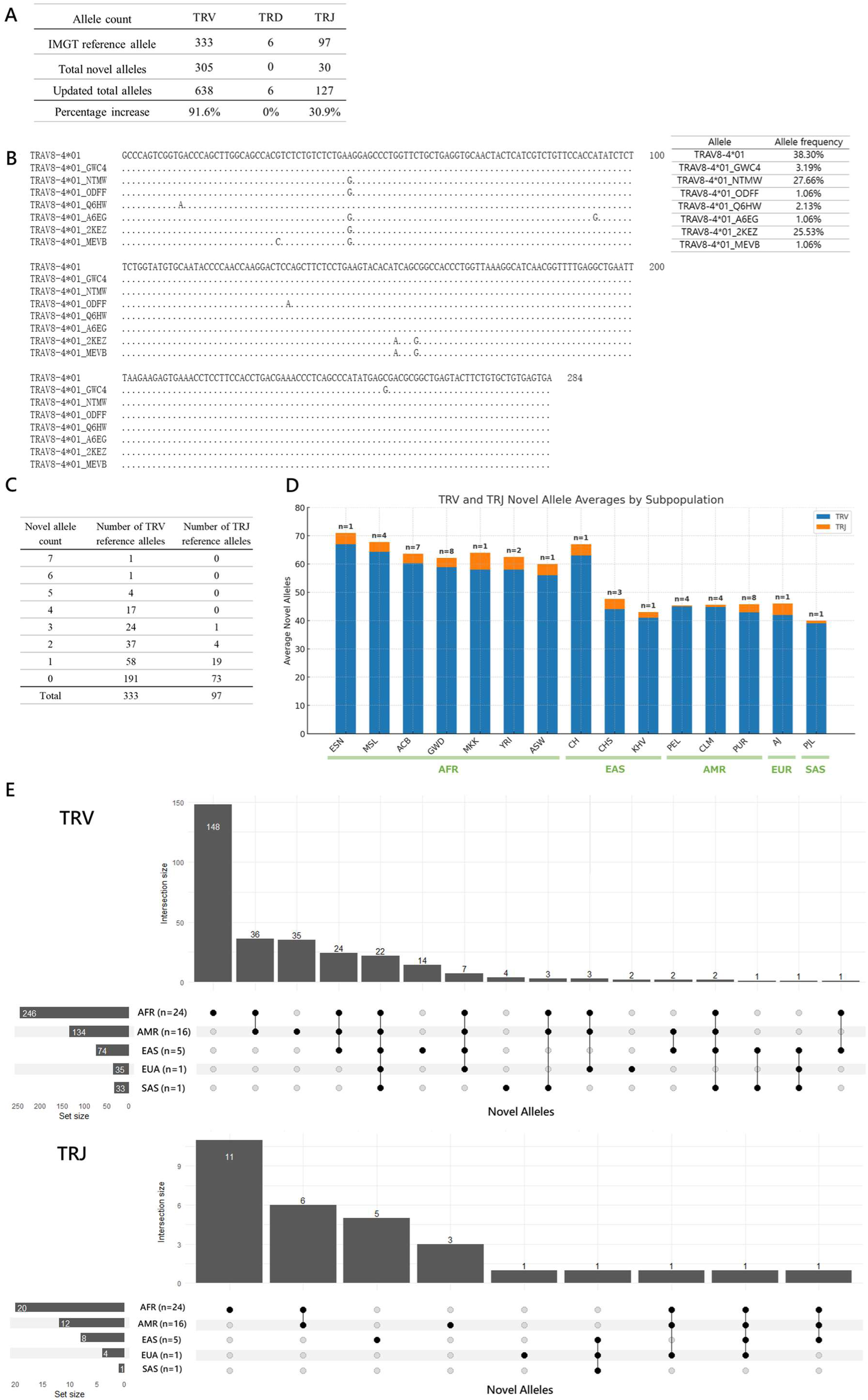
Expansion and characterization of the TR allele database. (A) Updated TR allele counts and percentage increase. IMGT/GENE-DB reference^1, 20^: Version 3.1.41 (2024-08-01); Accessed on 2025-04-19. (B) Sequence alignment of TRAV8-4*01 and its seven novel alleles. Sequence alignment of the TRAV8-4*01 reference allele (top row) with its seven novel alleles. Dots indicate sequence identity with the reference allele, while nucleotide differences are highlighted at their respective positions. The right panel shows the relative allele frequencies observed in 47 HPRC samples (94 haplotypes), with TRAV8-4*01 comprising 38.30% of the total, and novel alleles such as TRAV8-4*01_GWC4 and TRAV8-4*01_NTMW accounting for 3.19% and 27.66%, respectively. (C) Distribution of TR reference alleles by novel allele count. “Novel allele count” refers to the number of novel alleles identified for each reference allele listed in the IMGT database, such as TRAV8-4*01, which has seven novel alleles. (D) Average counts of novel TRV and TRJ alleles across superpopulations. The bar chart illustrates the average counts of novel TRV (blue) and TRJ (orange) alleles in superpopulations and their respective subpopulations within the HPRC dataset. The numbers above each bar (n) indicate the sample size for each subpopulation. (E) Distribution of novel TRV and TRJ alleles across superpopulations (UpSet plot). UpSet plots depict the intersection of novel TRV (top) and TRJ (bottom) alleles across five superpopulations: AFR, AMR, EAS, EUR, and SAS. Each vertical bar represents the number of alleles shared by the indicated combination of superpopulations (shown below the plot), while horizontal bars indicate the total number of novel alleles identified in each superpopulation. The AFR population harbors the largest number of unique novel alleles for both TRV and TRJ, reflecting its greater genetic diversity compared to other populations.

Among the 333 TRV alleles recorded in the IMGT database, 42.6% have corresponding novel alleles identified in this study. Notably, TRAV8-4*01 had the highest number of novel alleles (n = 7) (Figure 4b), followed by TRGV3*01 (6 novel alleles), and TRAV15*01, TRAV31*01, TRBV5-8*01, and TRGV4*02 (each with 5 novel alleles). Likewise, 24.7% of the 97 IMGT TRJ reference alleles had novel alleles. TRAJ47*01 exhibited the highest number, with 3 novel alleles, while TRAJ17*01, TRAJ44*01, TRAJ52*01, and TRGJP2*01 each had 2 novel alleles. The remaining TRJ genes typically carry one or no novel alleles (Figure 4c).

The IMGT database classifies allele functionality into three categories: functional, open reading frame (ORF), or pseudogene.^32, 33, 34^ Different alleles of the same gene may differ in functionality; for example, TRBV7-3*01 is classified as functional, whereas TRBV7-3*02 is an ORF. In this study, we examined the functional classification of the closest IMGT reference allele for each novel allele. The majority of these closest reference alleles were functional— 65.1% of TRV and 93.3% of TRJ alleles. ORFs accounted for 5.6% of TRV and 6.7% of TRJ alleles, while pseudogenes comprised 29.6% of TRV alleles. A more detailed analysis of functionality of these novel alleles—including transitions to ORF or pseudogene status—is presented later in Subsection 7.

Most novel alleles were frequently observed across the 47 HPRC subjects. Among the 305 novel TRV alleles, a total of 2466 instances were observed across 94 haplotypes from the 47 HPRC subjects. The most prevalent novel allele, TRAV35*01_QREI, was found in 66 of the 94 haplotypes. Overall, 10 novel TRV alleles were present in over 47 haplotypes, corresponding to allele frequencies above 50%. Among the 30 novel TRJ alleles, 135 occurrences were detected across the 94 haplotypes, with TRDJ4*01_G4F3 being the most common, observed in 33 haplotypes. Three novel TRJ alleles were identified in more than 10 haplotypes, indicating allele frequencies exceeding 10% (Supplementary Figure 4).

Finally, we examined the distribution of novel alleles at both the individual and population levels. At the individual level, the number of novel TRV alleles ranged from 33 to 77, and novel TRJ alleles from 0 to 6 (Supplementary Table 2). At the population level, the 47 HPRC subjects were categorized into five superpopulations: AFR (African), AMR (Admixed American), EAS (East Asian), EUR (European), and SAS (South Asian). AFR subjects exhibited the highest average novel allele count—60.3 for TRV and 3.6 for TRJ—followed by EAS with 47.2 TRV and 3.4 TRJ alleles on average (Figure 4d; Supplementary Table 2).

Figure 4e illustrates the shared and population-specific novel alleles across superpopulations. The AFR superpopulation exhibited the highest number of population-specific alleles, with 148 unique TRV and 11 unique TRJ alleles, reflecting greater genetic diversity in African populations. In contrast, EUR and SAS harbored fewer unique alleles than AFR, AMR, and EAS. However, these findings should be interpreted with caution, given the limited sample size and unequal subject numbers across different populations. A comprehensive summary of allele distributions and frequencies across populations is provided in Supplementary Table 2.

### 5. Establishment of a comprehensive flanking sequence database for TR alleles

In addition to variants within the core coding regions of TR genes, their flanking sequences also exhibit considerable variability. To systematically analyze this variability, we collected all available flanking sequences from the 47 HPRC subjects and grouped identical sequences, with each group representing a distinct haplotype. The gAIRR-annotate pipeline initially extracted 200 bp of flanking sequences from both the 5’ and 3’ ends of each allele. However, we observed that microsatellites within these 200 bp regions could interfere with downstream analyses. To avoid these confounding effects, we reduced the extracted flanking sequence length to 50 bp. Both 200 bp and 50 bp flanking sequences were grouped by sequence identity, with detailed information provided in Supplementary Table 3.

To facilitate consistent labeling and ease of identification, distinct flanking sequence groups for the same allele were named using the format: allele_name&flankX. For example, the allele TRAV33*01 was found to have four distinct flanking sequence groups, which were then labeled as TRAV33*01&flank1 through TRAV33*01&flank4.

Analysis of the 50 bp flanking sequences within the HPRC cohort revealed that V could exhibit up to five distinct flanking sequence groups, J alleles up to four, and D alleles up to two. As illustrative examples, TRBV12-4*01 possessed five distinct flanking sequence groups and TRBJ2-5*01 possessed four (Figure 5a), underscoring the sequence diversity present even beyond the core coding regions.

**Figure 5.**
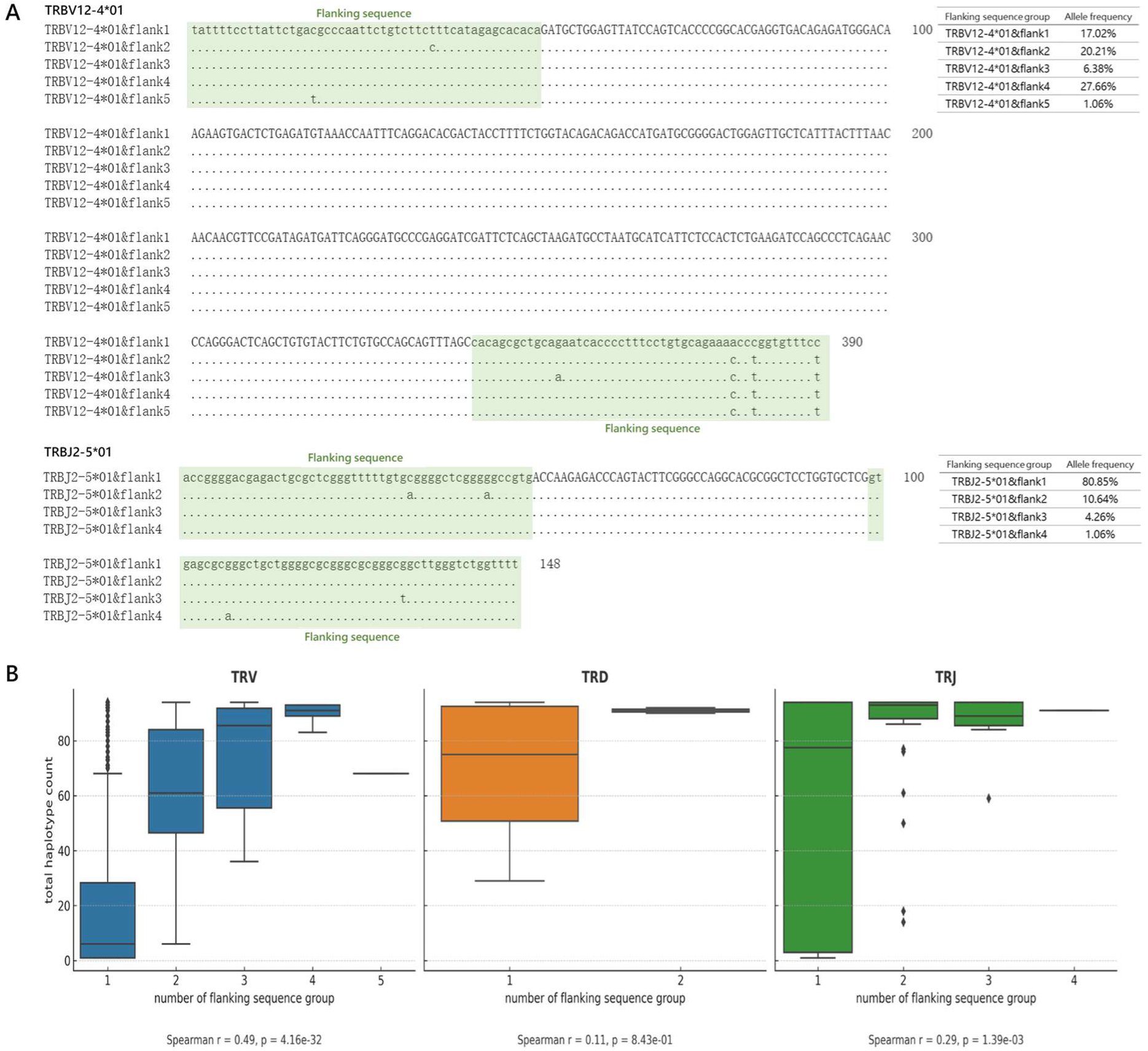
Flanking sequence variation and haplotype correlation. (A) Variation in flanking sequences for TRBV12-4*01 and TRBJ2-5*01 alleles. The green-shaded regions indicate flanking sequences (shown in lowercase), with 50 bp shown on each side of the core allele. TRBV12-4*01 exhibits five distinct flanking sequence groups, and TRBJ2-5*01 exhibits four. Each group represents a unique flanking variation identified across haplotypes within the HPRC cohort. Sequence identities are represented by dots (“.”). The right panels summarize the frequency of each flanking group across 94 haplotypes. For TRBV12-4*01, the most common flanking group is “flank4” (27.66%), while for TRBJ2-5*01, “flank1” predominates (80.85%). (B) Correlation between haplotype counts and flanking sequence groups for V, D, and J alleles. The boxplots illustrate the relationship between the number of distinct flanking sequence groups and total haplotype counts for each TR gene type (TRV, TRD, TRJ), based on 94 haplotypes from 47 HPRC individuals. The x-axis represents the number of flanking sequence groups observed for each allele, and the y-axis indicates the number of haplotypes carrying that allele. A positive correlation is observed between the number of flanking groups and haplotype count for TRV (Spearman’s r = 0.49, p = 4.16 × 10⁻³²) and TRJ (r = 0.29, p = 1.39 × 10⁻³), suggesting that alleles with broader representation tend to exhibit greater flanking diversity. No significant correlation was found for TRD alleles (r = 0.11, p = 0.843).

Further investigation revealed that some alleles exhibited more flanking sequence groups, demonstrating a higher degree of variability. We examined whether this variability was influenced by the prevalence of each allele in the population. Specifically, we performed analysis between the number of haplotypes carrying a given allele and the number of associated flanking sequence groups. The analysis revealed a positive correlation, indicating that alleles found in more individuals tended to exhibit greater flanking sequence diversity (Figure 5b). This result suggests that the observed variability may partly reflect broader population-level variation rather than allele-intrinsic hypervariability.

### 6. Characterization of RSS variability within flanking sequences

V(D)J recombination is mediated by RSS, which are located adjacent to each gene segment. An RSS consists of conserved heptamer and nonamer motifs separated by a spacer of either 12 or 23 bp.^5, 35^ As described previously, flanking sequences for each allele were grouped based on sequence similarity. RSS sites, characterized by typical lengths of 28 or 39 bp, reside immediately adjacent to gene segments, and the 50-bp flanking sequences extracted in our analysis were sufficient to encompass these RSS regions.

Based on the known RSS locations for different TR gene segments, we analyzed the 3’ flanking sequences of TRV alleles, the 5’ flanking sequences of TRJ alleles, and both ends for TRD alleles. These flanking sequences were evaluated using the RSSite tool for RSS prediction.^36^ According to known RSS features, expected RSS positions within the flanking sequences were defined as follows: positions 1-39 for the 3’ flanking sequence of TRV alleles, positions -28 to -1 for the 5’ flanking sequence for TRJ alleles, and positions 1-39 for the 3’ RSS and -28 to -1 for the 5’ RSS of TRD alleles (Figure 6a). Full prediction results are summarized in Supplementary Table 4.

**Figure 6.**
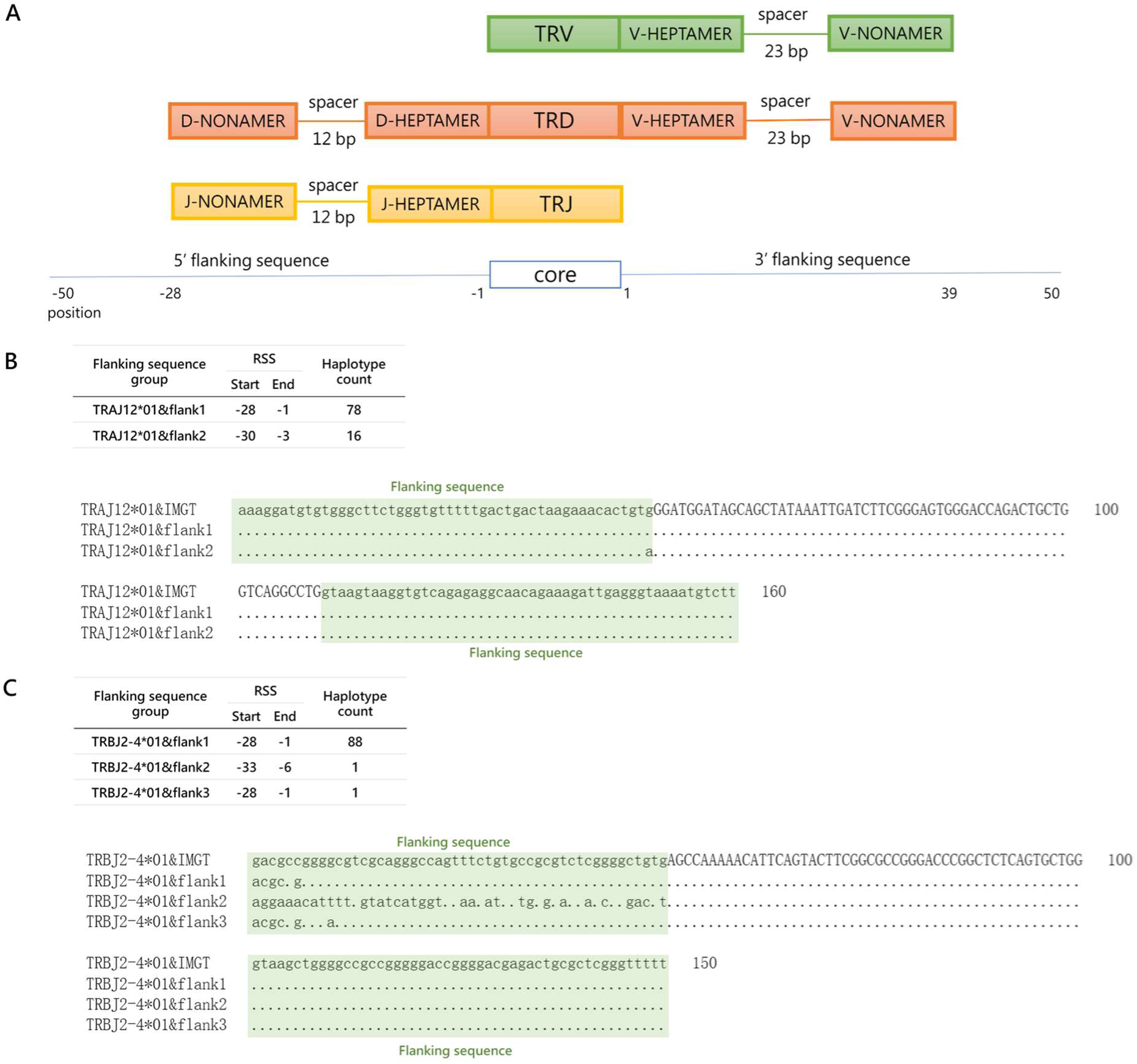
Further analysis of RSS in the flanking sequences. (A) Schematic representation of RSS in TRV, TRJ, and TRD genes. This schematic illustrates the RSSs adjacent to TRV, TRJ, and TRD genes. Each RSS consists of a conserved heptamer and nonamer motif, separated by a spacer region of 12 or 23 bp. The green boxes represent TRV gene, the yellow boxes represent TRJ, and the orange boxes represent TRD. The RSS positions within the 50 bp flanking sequences are expected as follows: for TRV alleles, the RSS is located between positions 1 – 39 in the 3’ flanking sequence; for TRJ alleles, it is located between positions -28 – -1 in the 5’ flanking sequence; and for TRD alleles, the 5’ RSS is positioned between 1 – 39, while the 3’ RSS is located between -28 – -1. (B) Identification of an unanticipated RSS in the flanking sequence of TRJ12*01. Flanking sequence comparison of TRJ12*01 reveals distinct RSS positioning patterns: one group (flank2) displays a shifted RSS site extending beyond the typical -28 to -1 region, with the -1 position nucleotide changing from G to A. The sequence labeled TRJ12*01&IMGT represents the flanking region obtained directly from the IMGT database, used here as a reference for comparison. (C) Identification of an unanticipated RSS in the flanking sequence of TRBJ2-4*01. Flanking sequence variation in TRBJ2-4*01 reveals an alternative RSS site (flank2) shifted upstream to positions -33 to -6. This shift is accompanied by extensive sequence divergence in the RSS region. The sequence labeled TRBJ2-4*01&IMGT represents the flanking region obtained directly from the IMGT database, used here as a reference for comparison.

While most RSS motifs were identified within their expected positional ranges, some exceptions were observed. For example, TRAV11*01 and TRAV14-1*01 exhibited RSS sites at positions 3-41, deviating from the expected locations. Both alleles are classified as pseudogenes by IMGT due to frameshift mutations in their V-REGION,^32, 37^ suggesting that aberrant RSS positioning may reflect or contribute to disrupted gene function.

In addition to alleles with inherently abnormal RSS positioning, we also identified several cases in which flanking sequence variations caused shifts in RSS location. For instance, TRAJ12*01 had two distinct flanking sequence groups. In TRAJ12*01&flank1, the RSS was located at the expected positions (-28 to -1), whereas in TRAJ12*01&flank2, the RSS shifted to positions -30 to -3 (Figure 6b). Similarly, TRBJ2-4*01 had three distinct flanking sequence groups, with the TRBJ2-4*01&flank2 group also exhibiting a shifted RSS. Sequence alignment revealed considerable flanking sequence divergence in this case, which was uniquely observed in the paternal haplotype of subject HG01978. Further investigation revealed a deletion of four genes—*TRBJ2-1*, *TRBJ2-2*, *TRBJ2-2P*, and *TRBJ2-3*—within this haplotype, likely contributing to the altered flanking sequence and RSS shift (Figure 6c).

These findings demonstrate that the same allele may exhibit distinct RSS architecture depending on flanking sequence context, potentially influencing recombination efficiency. Our results underscore the importance of precisely characterizing flanking sequence diversity—not only for accurate allele annotation but also for understanding functional implications in V(D)J recombination.

### 7. Evaluation of functional classifications and transitions among novel TR alleles

To assess the biological relevance of the novel TR alleles, we evaluated their predicted functionality using Digger,^38^ a tool that annotates TR alleles based on features such as the presence of intact open reading frames, conserved residues (e.g., cysteine bridges), and the absence of frameshift variants or premature stop codons. Our analysis focused on the 226 novel alleles whose closest reference alleles in the IMGT database were classified as functional— 198 TRV and 28 TRJ alleles.

Among these, the majority of novel alleles (86.9% of the TRV and 85.7% of the TRJ alleles) retained their status as functional. However, 13 TRV alleles were reclassified from functional to ORF. Additionally, 12 TRV alleles and 1 TRJ allele transitioned to pseudogene status, and 4 alleles could not be definitively classified (NA) (Table 1).

**Table 1.** Functional status changes in novel variant alleles. This table summarizes the predicted functional status of 226 novel TR alleles whose closest IMGT reference alleles were annotated as functional. Functional annotations were performed using Digger,^38^ based on features such as intact open reading frames, conserved residues, and the absence of frameshift or nonsense mutations.

| Functional status | TRV | TRJ |
| --- | --- | --- |
| Functional | 172 | 24 |
| ORF | 13 | 0 |
| Pseudogene | 12 | 1 |
| Not classified (NA) | 1 | 3 |
| Total | 198 | 28 |

A representative example is the TRAV10*01 locus, where three novel variants were identified: TRAV10*01_IM77, TRAV10*01_DOMV, and TRAV10*01_S23D. While TRAV10*01_IM77 and TRAV10*01_S23D preserved key functional motifs, TRAV10*01_DOMV was reclassified as an ORF due to the absence of a critical cysteine residue.

These findings suggest that a subset of novel alleles may have a substantial impact on TR structure and function. Given that germline TR alleles have been previously associated with conditions such as autoimmune disease,^14, 15^ and responses to immunotherapy,^16^ underscoring the need for future studies to investigate the immunological consequences of these novel alleles. These results highlight the value of incorporating high-resolution germline annotations into future immunogenomic and clinical association studies.

## Discussion

In this study, we leveraged high-quality phased diploid genome assemblies from HPRC^25^ to comprehensively characterize the germline gene set encoding TR. Using the gAIRR Suite,^21^ a modular pipeline composed of gAIRR-annotate, gAIRR-seq, and gAIRR-call, we identified 335 novel TR alleles, including 305 TRV and 30 TRJ alleles, representing 91.6% and 30.9% increases over existing IMGT records, respectively. Importantly, all novel alleles were crosschecked using two orthogonal approaches: genome assembly-based annotation and capture-based short-read sequencing followed by read-based genotyping. This dual validation strategy, combined with flanking sequence integration, provided strong support for the authenticity and accuracy of the identified alleles.

In addition to core region expansion, we established a comprehensive database of upstream and downstream flanking sequences—including RSS—for all detected alleles. Our flanking sequence analysis revealed previously unrecognized sequence diversity, including RSS variability that may influence recombination efficiency.^5, 6, 7^ Moreover, by incorporating flanking sequence identity into allele classification, we were able to identify additional novel alleles that would otherwise be misclassified or missed entirely by core-sequence-only approaches. Collectively, our study substantially expands the known TR germline landscape and provides valuable reference resources for immunogenomics research.

Given the success of this approach, we advocate for its expansion to larger datasets. Future applications to the ongoing project of Year-2 HPRC cohort,^25^ which includes a broader range of global populations and higher sample size, would be especially valuable. Furthermore, other high-resolution genome projects, such as the All of Us Research Program (NIH),^39^ and projects supported by the Telomere-to-Telomere (T2T) Consortium,^40^ offer promising opportunities— provided they produce long-read whole-genome sequencing (WGS) data and make high-quality personal genome assemblies and matched genomic DNA samples publicly available. While our current analysis demonstrated significant diversity, population coverage remains incomplete; notably, East and Southeast Asian representation is still limited in the HPRC,^25^ which underscores the need for broader sampling to achieve equitable immunogenomic discovery.

Our approach is particularly well-suited for TR allele discovery, but less ideal for IG genes, due to the use of Epstein–Barr virus (EBV)-transformed B lymphocyte cell lines in many reference datasets. Such transformation introduces somatic V(D)J recombination events in IG loci, thereby complicating the inference of true germline configurations.^41, 42, 43^ Alternative strategies, including single-cell VDJ mapping or using untransformed genomic DNA from non-lymphoid tissues, may be required for IG-focused germline analysis.

A critical challenge facing the field is the lack of an efficient, community-accepted framework for naming, curating, and integrating novel gAIRR alleles into standardized reference sets. While the AIRR Community and recent efforts such as those by Collins et al. (2024)^27^ and Lees et al. (2023)^28^ have established curation standards and schemas for IG genes, the process of official allele approval and deposition into databases such as IMGT^1^ remains labor-intensive and slow. The absence of automated or scalable submission pathways hinders timely updates and broader adoption of gAIRR discoveries. Continued development of tools like IgLabel^28^ and collaboration with community-led repositories will be essential for streamlining this process.

Importantly, gAIRR data serves as the reference backbone for profiling the exprAIRR, obtained through AIRR-seq technologies. Inaccurate or incomplete gAIRR references can directly compromise exprAIRR analysis, leading to erroneous clonotype assignments and interpretation.^9^ ^10^ Furthermore, the value of gAIRR data extends beyond serving as the references for exprAIRR profiling. Several germline TR alleles have been associated with diverse immune-related phenotypes and diseases, including, autoimmune disorders,^14, 15^ vaccine responsiveness.^11^ As such, population-scale gAIRR allele maps offer a foundation for genotype-phenotype association studies that can uncover new immunogenetic mechanisms.

Looking ahead, integrated analyses that combine gAIRR genotyping, bulk or single-cell exprAIRR profiling, and HLA allele resolution represent a powerful strategy for dissecting immune variation. Such multi-modal approaches could elucidate the interplay between germline receptor architecture, transcriptional expression, and peptide presentation in shaping immune responses to infection, autoimmunity, cancer, and therapeutic interventions. With the expansion of reference resources, scalable sequencing technologies, and computational infrastructure, we envision the integrated gAIRR + exprAIRR + HLA framework as a cornerstone of precision immunology.

### Limitations of the study

While this study provides a robust framework for identifying and validating novel TR alleles, several limitations remain. First, our analysis relied on genomic data from 47 individuals representing five superpopulations. Although this sample size was sufficient for detecting common TR variants, it may have limited our ability to identify rarer variants. Additionally, the subjects included in the HPRC dataset were predominantly African, potentially biasing allele discovery toward variants prevalent in African populations and thus limiting the representation of global TR allele diversity. Lastly, discrepancies between the gAIRR-annotate and gAIRR-call results, such as errors in genome assemblies observed for specific subjects, highlight ongoing challenges and the necessity for further refinement of genome assembly methods and annotation pipelines.

## Resource availability

### Lead contact

Further information and requests for resources and reagents should be directed to and will be fulfilled by the lead contact, Pei-Lung Chen.

#### Data and code availability

This paper does not report any original code. However, the primary analysis was conducted using the gAIRR Suite, and its source code is available at https://github.com/maojanlin/gAIRRsuite. The sequences of the 335 novel TR alleles identified in this study are publicly accessible at https://zenodo.org/records/15069123. Additionally, the comprehensive flanking sequence database, which includes both novel and reference alleles, is available at https://zenodo.org/records/15069181.

## Acknowledgments

We would like to acknowledge the National Human Genome Research Institute (NHGRI) for funding the following grants supporting the creation of the human pangenome reference: 1U41HG010972, 1U01HG010971, 1U01HG013760, 1U01HG010961, 1U01HG010973, 1U01HG013748, 1U01HG013744, 1U01HG013755, 1U01HG010963, and the Human Pangenome Reference Consortium (https://humanpangenome.org/). We thank the Coriell Institute for providing the cell lines used in this study. DNA samples were obtained from the NHGRI Sample Repository for Human Genetic Research, maintained by the Coriell Institute for Medical Research. We would like to thank the National Center for High-performance Computing (NCHC) of the National Institutes of Applied Research (NIAR) of Taiwan for providing computational and storage resources. We would like to thank the following grants: UN111-061, 111-2314-B-002-243-MY3, 112-TMU211, 112L895202, 113L892902, 114L891102, NSTC112-2622-B-002-008-, and VN113-14.

## Author contributions

P.L. Chen conceived the initial project. I.H. Chiu, S.K. Lai, and C.Y. Chen conducted the experiments. Y.H. Yang, C.Y. Yao, M. J. Lin, K.T. Huang, and Y.H. Lin performed the data analysis. Y.H. Yang drafted the manuscript. Y.C. Yang, C.L. Hsu, J.S. Hsu, C.Y. Chen, and P.L. Chen jointly supervised the project.

## Declaration of interests

The authors declare no competing interests.

## STAR★Methods

### Key resources table

| REAGENT or RESOURCES | SOURCE | IDENTIFIER |
| --- | --- | --- |
| **Deposited data** |  |  |
| Assemblies | HPRC<sup>25</sup> | https://github.com/human-pangenomics/HPP_Year1_Assemblies |
| TR reference alleles | IMGT<sup>1, 20</sup> | https://www.imgt.org/IMGTrepertoire/ |
| **Software and algorithms** |  |  |
| gAIRR Suite | M.J. Lin et al.<sup>21</sup> | https://github.com/maojanlin/gAIRRsuite |

## Methods

### Reference materials

We used assemblies from 47 subjects provided by the HPRC^25^ (https://github.com/human-pangenomics/HPP_Year1_Data_Freeze_v1.0). Genomic DNA reference materials for these subjects were obtained from the Coriell Institute (https://www.coriell.org). The genomic DNA samples were extracted from EBV-transformed B lymphocytes. As a reference for TR gene annotation, we used the IMGT/GENE-DB reference database^1, 20^ (Version 3.1.41, released on 2024-08-01), accessed on 2025-04-19.

### gAIRR Suite pipeline overview

Assemblies were analyzed using gAIRR-annotate to identify TR alleles. DNA obtained from the Coriell Institute underwent library preparation, target region capture, and sequencing, a process referred to as gAIRR-seq. The FASTQ files generated from gAIRR-seq were processed using gAIRR-call to identify TR alleles. For detailed methods, please refer to our previously published paper.^21^

### Renaming of novel alleles using IgLabel and updating gAIRR Suite

The gAIRR-annotate module identifies alleles from genome assemblies by aligning sequences to reference alleles. If variants are detected, they are labeled as novel alleles based on the closest matching reference allele. Novel alleles identified by gAIRR-annotate are initially marked with the suffix /novel.

To systematically manage these novel alleles, we used IgLabel.^28^ The sequences of novel alleles identified from the 47 HPRC assemblies were collected and imported into IgLabel. IgLabel analyzes the sequence differences and assigns a unique four-character code to each allele, which is appended to the allele label for differentiation.

After renaming the novel alleles, they were incorporated into the gAIRR Suite materials. Finally, gAIRR-annotate was rerun to ensure all novel alleles were accurately named and integrated into the pipeline.

### Visualization of results with IGV

FASTQ files from gAIRR-seq were aligned to the reference FASTA using BWA (v0.7.17-r1188) [^44^], producing SAM files. These SAM files were then converted to BAM files using SAMtools (v1.19.1).^45^ Finally, the sequencing results were visualized and validated using IGV (v2.10.3).^46^ For visualization, a Mapping Quality (MPQ) threshold of 20 and an Alignment Score threshold of 300 were applied.

### Using the Digger to identify the functionality of novel alleles

We analyzed the assemblies of 47 HPRC subjects using Digger^38^ to identify alleles and gather information on contigs, positions, and their predicted functionality. We then compared these results with those from gAIRR-annotate, focusing on the contigs and positions of each novel allele. This comparison allowed us to determine the functionality of each novel allele as predicted by Digger.

### Flanking sequence collection and grouping

Flanking sequences for each allele were collected using the gAIRR-annotate tool, which processes samples to extract the flanking sequences of all identified alleles. This analysis was performed on all 47 HPRC subjects, resulting in a comprehensive collection of flanking sequences for each allele. Variants were observed in the flanking sequences of the same allele across different samples; therefore, sequences with identical flanking regions were grouped together to define unique haplotypes. While the original design of gAIRR-annotate extracted 200 bp flanking sequences, we observed interference from microsatellites in these regions. To address this issue, we also extracted 50 bp flanking sequences. Both 50 bp and 200 bp sequences were subsequently grouped to account for flanking sequence variability.

### RSS site prediction using RSSite

To predict RSS sites, we utilized the RSSite tool.^36^ RSS sites are located on either side of the gene and have conserved lengths of 28 bp or 39 bp (Figure 8). The 50 bp flanking sequences extracted in our analysis were sufficient to encompass these RSS regions. For this prediction, we used the curated 50 bp flanking sequences grouped for each allele. Specifically, we used the 5’ end flanking sequences for TRV alleles, the 3’ end for TRJ alleles, and both ends for TRD alleles. These 50 bp flanking sequences were input into the RSSite web tool to predict the exact location of RSS sites for each allele.

## Supplemental information

Document S1. Figures S1–S4

Table S1. Sequences of novel alleles, related to Figure 4a, 4b and 4c.

Table S2. Distribution of novel alleles across individuals and populations, related to Figure 4d and 4e.

Table S3. Flanking sequence database, related to Figure 5. Table S4. Predicted RSS positions, related to Figure 6.

## Notes

https://zenodo.org/records/15069123

https://zenodo.org/records/15069181

### Competing Interest Statement

The authors have declared no competing interest.

