## Supplementary Figures and Tables for "Unmasking Human T Cell Receptor Germline Diversity: 335 Novel Alleles Identified in 47 Pangenome Reference Individuals Using the gAIRR Suite": TR_novel_supplementary_figure.pdf

Supplementary figure 1: Analysis of mismatches in novel allele validation. Can be divided into three parts: 1-1, 1-2 and 1-3.

a.

```
M33233|TRBV11-1*01|Homo       gaagctgaagttgcccagtcccccagatataagattacagagaaaagccaggctgtggct    60
new|TRBV11-1*01_B6R4|Homo     gaagctgaagttgcccagtcccccagatataagattacagagaaaagccaggctgtggct    60
                              ************************************************************

M33233|TRBV11-1*01|Homo       ttttggtgtgatcctatttctggccatgctaccctttactggtacc-ggcagatcctggg    119
new|TRBV11-1*01_B6R4|Homo     ttttggtgtgatcctatttctggccatgctaccctttactggtacccggcagatcctggg    120
                              ********************************************** *************

M33233|TRBV11-1*01|Homo       acagggcccggagcttctggttcaatttcaggatgagagtgtagtagatgattcacagtt    179
new|TRBV11-1*01_B6R4|Homo     acagggcccggagcttctggttcaatttcaggatgagagtgtagtagatgattcacagtt    180
                              ************************************************************

M33233|TRBV11-1*01|Homo       gcctaaggatcgattttctgcagagaggctcaaaggagtagactccactctcaagatcca    239
new|TRBV11-1*01_B6R4|Homo     gcctaaggatcgattttctgcagagaggctcaaaggagtagactccactctcaagatcca    240
                              ************************************************************

M33233|TRBV11-1*01|Homo       gcctgcagagcttggggactcggccatgtatctctgtgccagcagcttagc    290
new|TRBV11-1*01_B6R4|Homo     gcctgcagagcttggggactcggccatgtatctctgtgccagcagcttagc    291
                              ***************************************************
```

b.

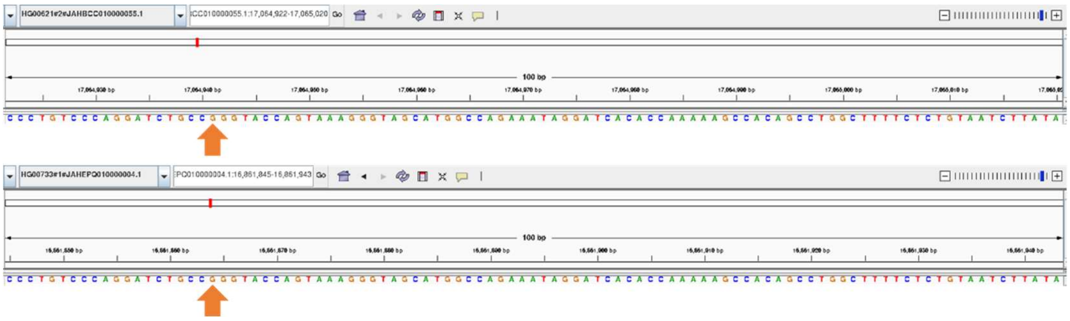

c.

TRBV11-1*01

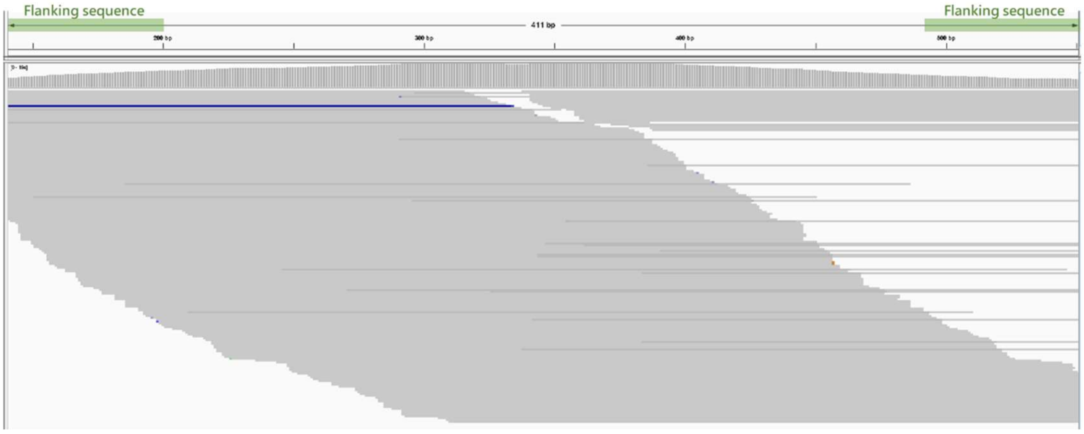


TRBV11-1*01_B6R4

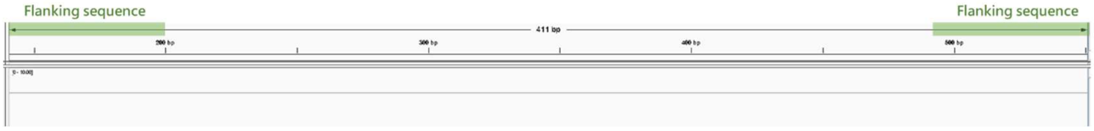

d.

HG00621

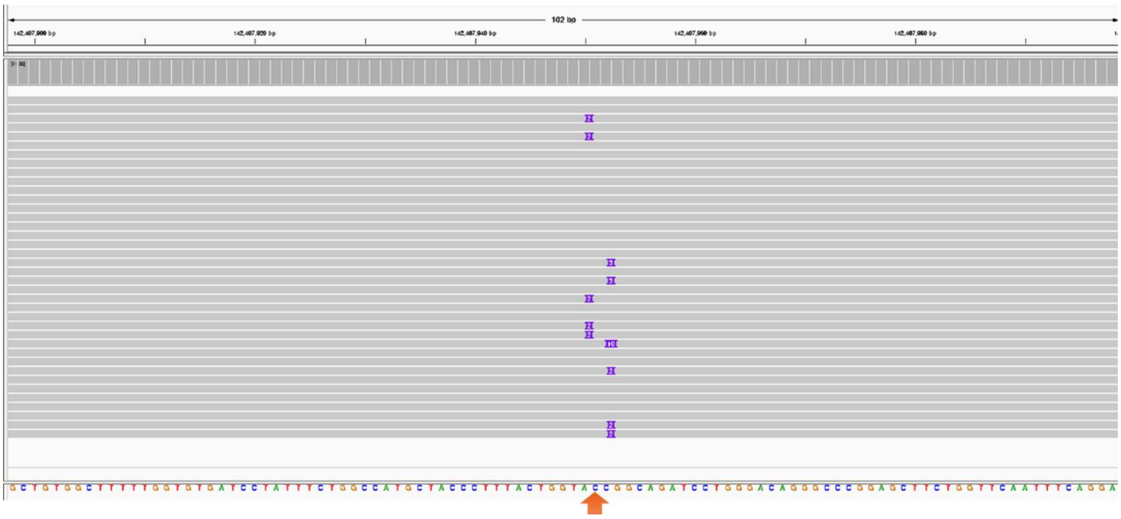

HG00733

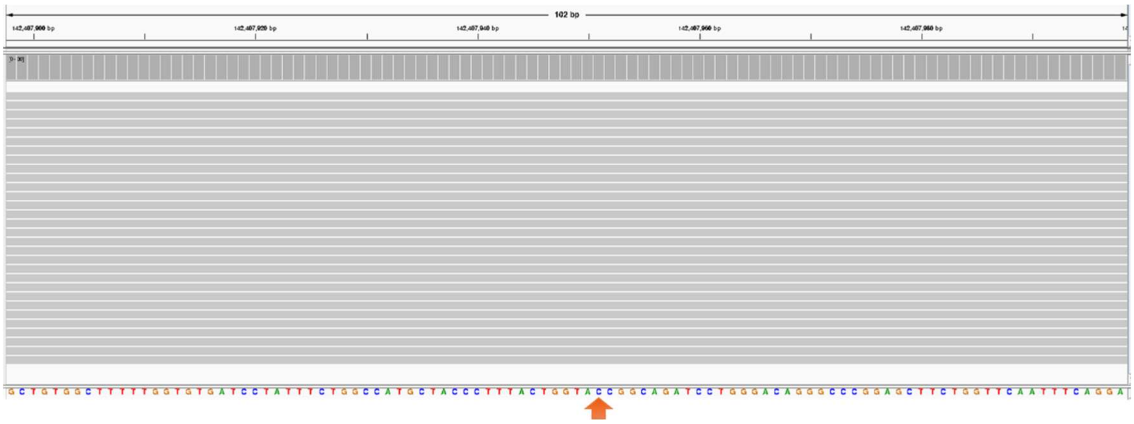

**Supplementary figure 1-1: Analysis of Mismatches in TRBV11-1*01_B6R4.**

gAIRR-annotate identified that HG00621 and HG00733 possess the novel allele TRBV11-1*01_B6R4. However, gAIRR-call did not confirm this finding. Below is the detailed analysis:

(a) TRBV11-1*01_B6R4 is a novel allele derived from TRBV11-1*01, featuring an insertion of a "C" at position 106, changing a stretch of two consecutive "C"s into three.

(b) To verify this insertion, we used the positional information provided by gAIRR-annotate and visualized the sequences using IGV. The results clearly show that HG00621 and HG00733 both have three "G"s (due to the reverse strand orientation of the assembly sequences).

(c) Next, we aligned the gAIRR-seq fastq data to both TRBV11-1*01 and TRBV11-1*01_B6R4. The alignment results indicate that reads only map to TRBV11-1*01, and no reads align to TRBV11-1*01_B6R4.

(d) Finally, we downloaded HG00621_aligned_GRCh38_winnowmap.sorted.bam and HG00733_aligned_GRCh38_winnowmap.sorted.bam from the HPRC web resources. Upon examining the GRCh38 TRBV11-1 position, we observed that HiFi long reads from HG00621 support the insertion. However, the insertion is absent in HG00733, indicating an assembly error in HG00733.

a.

```
X74798|TRGV10*02|Homo        ttatcaaaagtggagcagttccagctatccatttccacggaagtcaagaaaagtattgac    60
new|TRGV10*02_FO2S|Homo      ttatcaaaagtggagcagttccagctatccatttccacggaagtcaagaaaagtattgac    60
                             ************************************************************

X74798|TRGV10*02|Homo        ataccttgcaagatatcgagcacaaggtttgaaacagatgtcattcactggtaccg-gca    119
new|TRGV10*02_FO2S|Homo      ataccttgcaagatatcgagcacaaggtttgaaacagatgtcattcactggtacccggca    120
                             ********************************************************  ***

X74798|TRGV10*02|Homo        gaaaccaaatcaggctttggagcacctgatctatattgtctcaacaaaatccgcagctcg    179
new|TRGV10*02_FO2S|Homo      gaaaccaaatcaggctttggagcacctgatctatattgtctcaacaaaatccgcagctcg    180
                             ************************************************************

X74798|TRGV10*02|Homo        acgcagcatgggtaagacaagcaacaaagtggaggcaagaaagaattctcaaactctcac    239
new|TRGV10*02_FO2S|Homo      acgcagcatgggtaagacaagcaacaaagtggaggcaagaaagaattctcaaactctcac    240
                             ************************************************************

X74798|TRGV10*02|Homo        ttcaatccttaccatcaagtccgtagagaaagaagacatggccgtttactactgtgctgc    299
new|TRGV10*02_FO2S|Homo      ttcaatccttaccatcaagtccgtagagaaagaagacatggccgtttactactgtgctgc    300
                             ************************************************************

X74798|TRGV10*02|Homo        gtgggattac 309
new|TRGV10*02_FO2S|Homo      gtgggattac 310
                             **********
```

b.

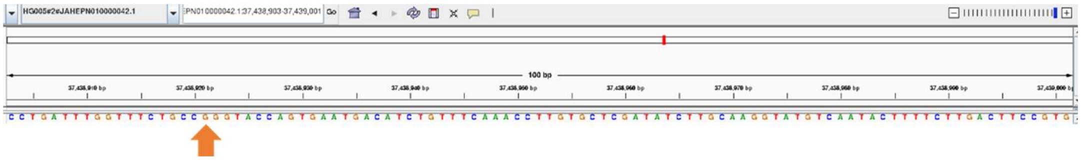

c.

TRGV10*02

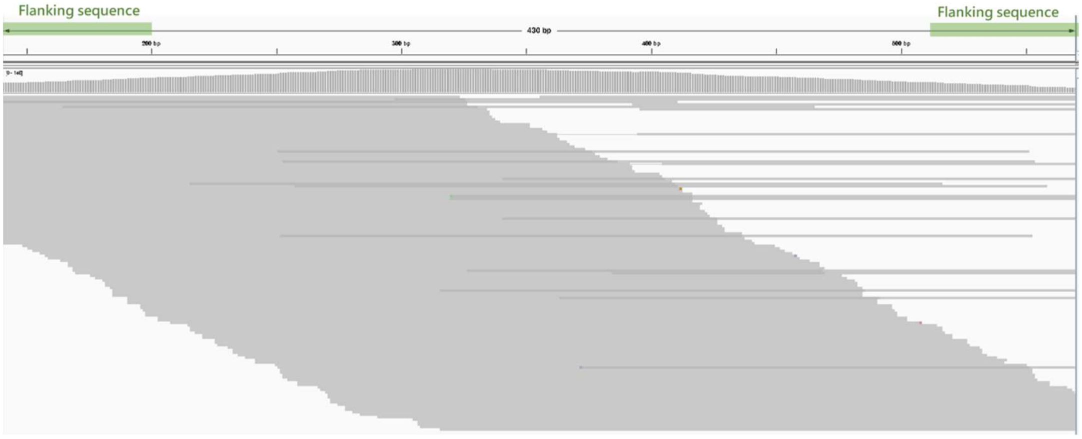


TRGV10*02_FO2S

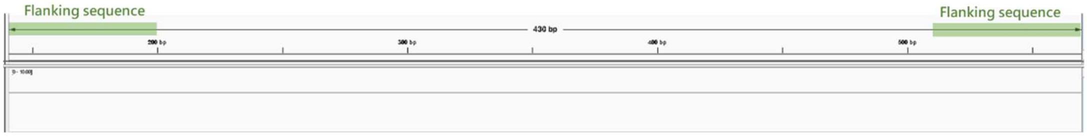

d.

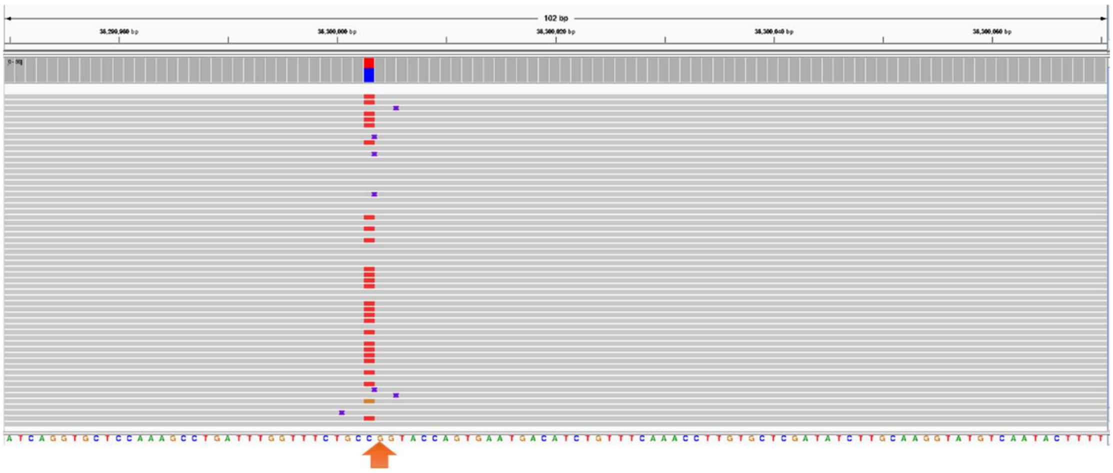

**Supplementary figure 1-2: Analysis of Mismatches in TRGV10*02_FO2S.**

gAIRR-annotate identified that HG005 possess the novel allele TRGV10*02_FO2S. However, gAIRR-call did not confirm this finding. Below is the detailed analysis:

(a) TRGV10*02_FO2S is a novel allele derived from TRGV10*02, featuring an insertion of a "C" at position 115, changing a stretch of two consecutive "C"s into three.

(b) To verify this insertion, we used the positional information provided by gAIRR-annotate and visualized the sequences using IGV. The results clearly show that HG005 have three "G"s (due to the reverse strand orientation of the assembly sequences).

(c) Next, we aligned the gAIRR-seq fastq data to both TRGV10*02 and TRGV10*02_FO2S. The alignment results indicate that reads only map to TRGV10*02, and no reads align to TRGV10*02_FO2S.

(d) Finally, we downloaded HG005_aligned_GRCh38_winnowmap.sorted.bam from the HPRC web resources. Upon examining the GRCh38 TRGV10*02 position, we observed that HiFi long reads from HG005 support the insertion.

a.

```
TRBJ2-7*01          ctcctacgagcagtacttcgggccgggcaccaggctcacggtcacag    47
TRBJ2-7*01_UZLC     TAACTACGAGCAGTACTTCGGGCCGGGCACCAGGCTCACGGTCACAG    47
                      *********************************************
```

b.

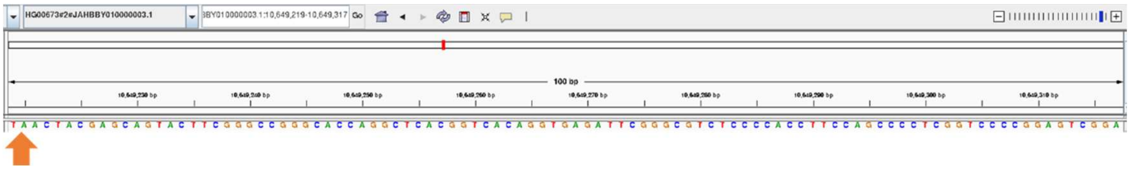

c.

TRBJ2-7*01

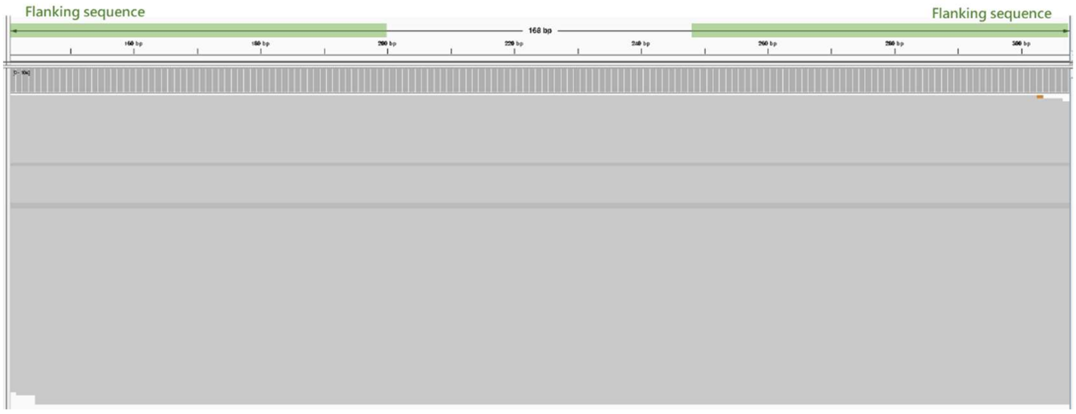


TRBJ2-7*01_UZLC

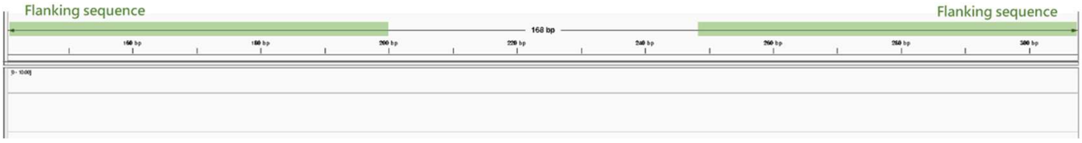


d.

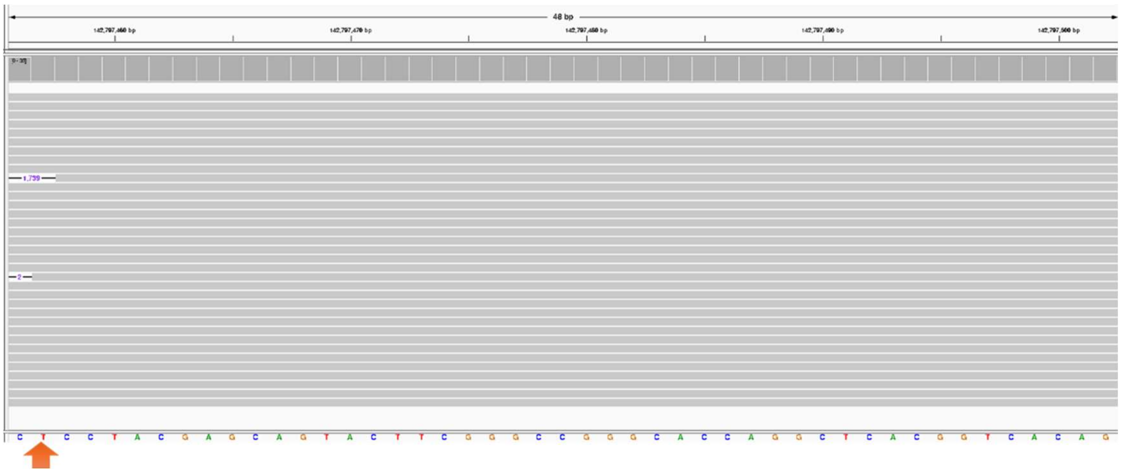


**Supplementary figure 1-3: Analysis of Mismatches in TRBJ2-7*01_UZLC.**

gAIRR-annotate identified that HG00673 possesses the novel allele TRBJ2-7*01_UZLC. However, gAIRR-call did not confirm this finding. Below is the detailed analysis:

(a) TRBJ2-7*01_UZLC is a novel allele derived from TRBJ2-7*01, where the sequence "CTC" at positions 1 to 3 is replaced by "TAA".

(b) To verify this substitution, we used the positional information provided by gAIRR-annotate and visualized the sequences using IGV. The results clearly show that HG00673 has the sequence "TAA" at this position.

(c) Next, we aligned the gAIRR-seq fastq data to both TRBJ2-7*01 and TRBJ2-7*01_UZLC. The alignment results indicate that reads only map to TRBJ2-7*01, and no reads align to TRBJ2-7*01_UZLC.

(d) Finally, we downloaded HG00673_aligned_GRCh38_winnowmap.sorted.bam from the HPRC web resources. Upon examining the GRCh38 TRBJ2-7*01 position, we observed that the HiFi long reads from HG00673 contain the sequence “CTC”, indicating that the “TAA” substitution is not supported and confirming an assembly error in HG00673.

a.

| Flanking sequence length | TRV allele count | TRD allele count | TRJ allele count |
|---|---|---|---|
| 0 bp (Left) | 27 | 0 | 0 |
| 1-49 bp (Left) | 18 | 1 | 10 |
| ≥50 bp (Left) | 288 | 5 | 87 |
| 0 bp (Right) | 61 | 0 | 1 |
| 1-49 bp (Right) | 31 | 0 | 14 |
| ≥50 bp (Right) | 241 | 6 | 82 |
| 0 bp on Both Sides | 15 | 0 | 0 |
| ≥50 bp on Both Sides | 219 | 5 | 82 |
| Ratio | 65.8% (219/333) | 83.3% (5/6) | 84.5% (82/97) |

b.

| Allele | Left | Right | Allele | Left | Right |
|---|---|---|---|---|---|
| TRAV1-1*02 | 50 | 50 | TRBV10-3*03 | 0 | 0 |
| TRAV12-2*02 | 50 | 50 | TRBV11-3*04 | 50 | 49 |
| TRAV12-2*03 | 0 | 50 | TRBV14*02 | 45 | 0 |
| TRAV14/DV4*03 | 50 | 50 | TRBV19*03 | 50 | 50 |
| TRAV21*02 | 50 | 0 | TRBV20-1*02 | 50 | 0 |
| TRAV23/DV6*04 | 0 | 0 | TRBV20-1*05 | 50 | 0 |
| TRAV35*02 | 50 | 0 | TRBV30*04 | 0 | 50 |
| TRAV36/DV7*04 | 50 | 32 | TRBV4-3*04 | 0 | 0 |
| TRAV38-1*03 | 34 | 50 | TRBV5-5*02 | 50 | 0 |
| TRAV6*02 | 50 | 0 | TRBV7-5*01 | 50 | 50 |
| TRAV6*03 | 0 | 0 | TRDV2*02 | 0 | 0 |
| TRAV8-4*03 | 50 | 0 | TRGV2*02 | 43 | 50 |
| TRAV9-2*03 | 50 | 0 | TRGV3*02 | 50 | 50 |
| TRBV2*02 | 0 | 0 | TRAJ47*02 | 50 | 35 |
| TRBV10-1*03 | 0 | 0 | | | |

**Supplementary figure 2: Completeness and availability of flanking sequences for TR alleles in the IMGT database**

(a) Completeness of flanking sequences for TR alleles in the IMGT database.

(b) Flanking sequence availability for 29 IMGT alleles corresponding to novel alleles

a.

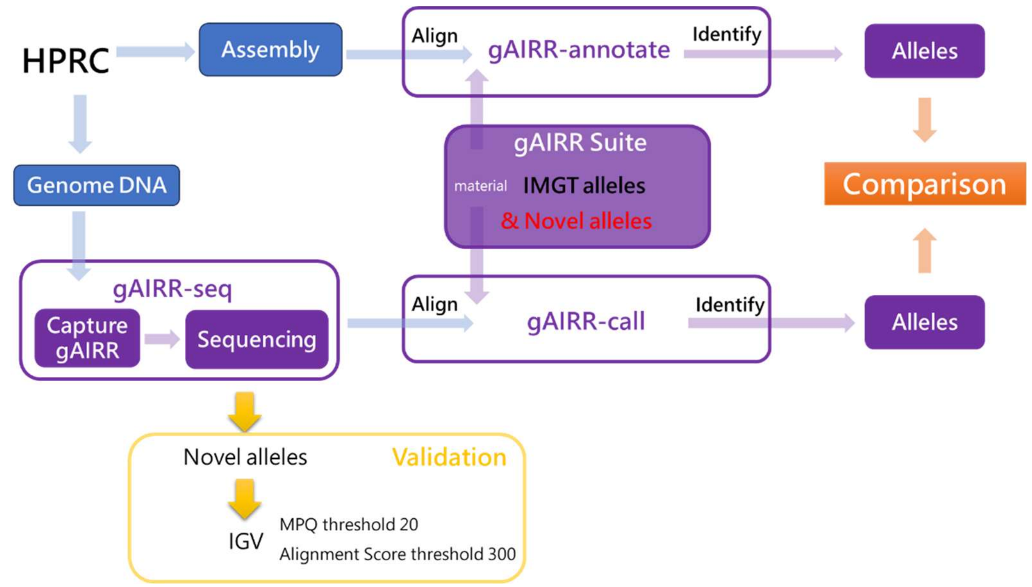


b.

TRAV8-4*01

```
TRAV8-4*01        GCCCAGTCGGTGACCCAGCTTGGCAGCCACGTCTCTGTCTCTGAAGGAGCCCTGGTTCTGCTGAGGTGCAACTACTCATCGTCTGTTCCACCATATCTCT   100
TRAV8-4*01_GWC4   ....................................................................................................
TRAV8-4*01_NTMW   ............................................G.......................................................
TRAV8-4*01_ODFF   ............................................G.......................................................
TRAV8-4*01_Q6HW   ...........A........................................................................................
TRAV8-4*01_A6EG   ............................................G...............................................G.......
TRAV8-4*01_2KEZ   ............................................G.......................................................
TRAV8-4*01_MEVB   ..............................C.............G.......................................................

TRAV8-4*01        TCTGGTATGTGCAATACCCCAACCAAGGACTCCAGCTTCTCCTGAAGTACACATCAGCGGCCACCCTGGTTAAAGGCATCAACGGTTTTGAGGCTGAATT   200
TRAV8-4*01_GWC4   ....................................................................................................
TRAV8-4*01_NTMW   ....................................................................................................
TRAV8-4*01_ODFF   ................................A...................................................................
TRAV8-4*01_Q6HW   ....................................................................................................
TRAV8-4*01_A6EG   ....................................................................................................
TRAV8-4*01_2KEZ   .....................................................A...G..........................................
TRAV8-4*01_MEVB   .....................................................A...G..........................................

TRAV8-4*01        TAAGAAGAGTGAAACCTCCTTCCACCTGACGAAACCCTCAGCCCATATGAGCGACGCGGCTGAGTACTTCTGTGCTGTGAGTGA   284
TRAV8-4*01_GWC4   ...................................................G................................
TRAV8-4*01_NTMW   ....................................................................................
TRAV8-4*01_ODFF   ....................................................................................
TRAV8-4*01_Q6HW   ....................................................................................
TRAV8-4*01_A6EG   ....................................................................................
TRAV8-4*01_2KEZ   ....................................................................................
TRAV8-4*01_MEVB   ....................................................................................
```

HG02572: TRAV8-4*01_2KEZ (homozygous)

TRAV8-4*01

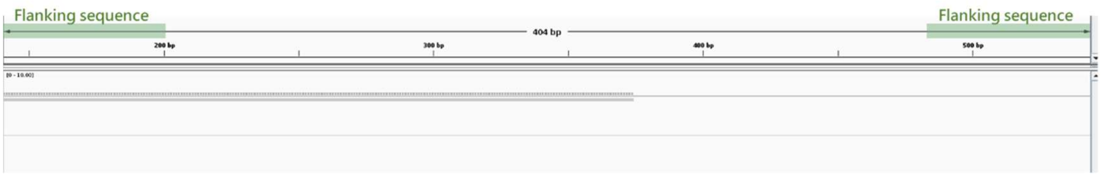

TRAV8-4*01_2KEZ

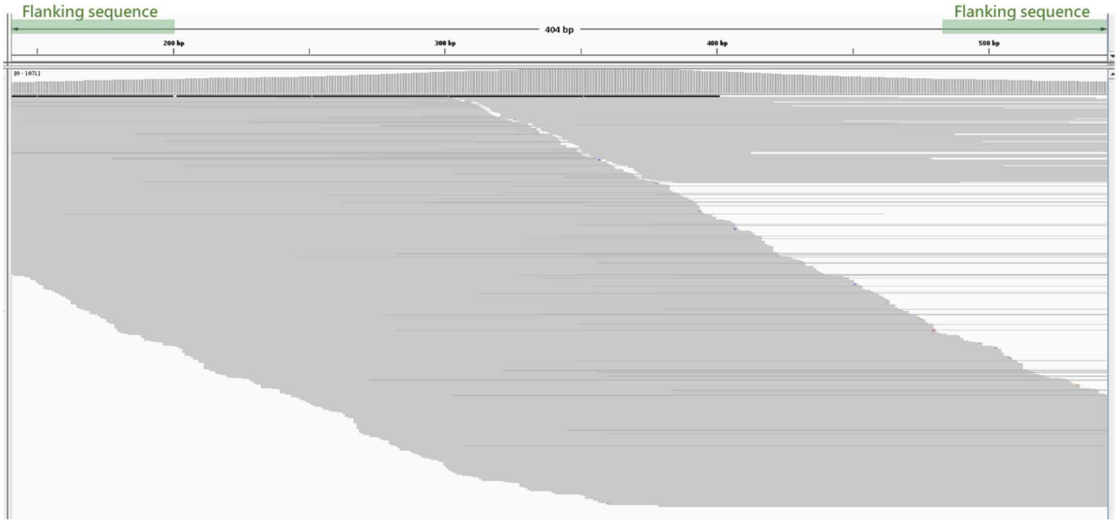


HG02257: TRAV8-4*01_A6EG & TRAV8-4*01_NTMW

TRAV8-4*01

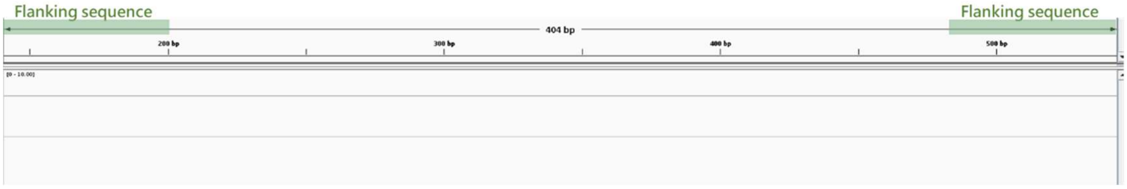


TRAV8-4*01_A6EG

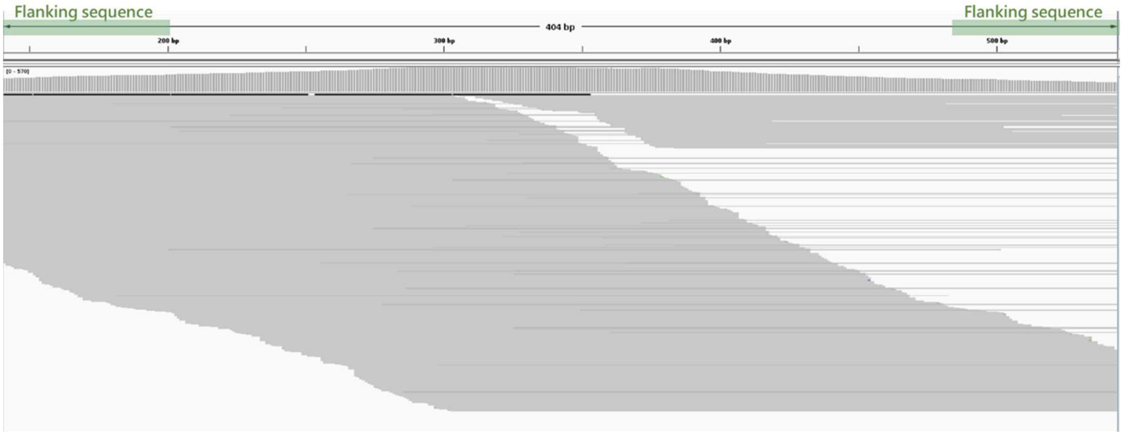


TRAV8-4*01_NTMW

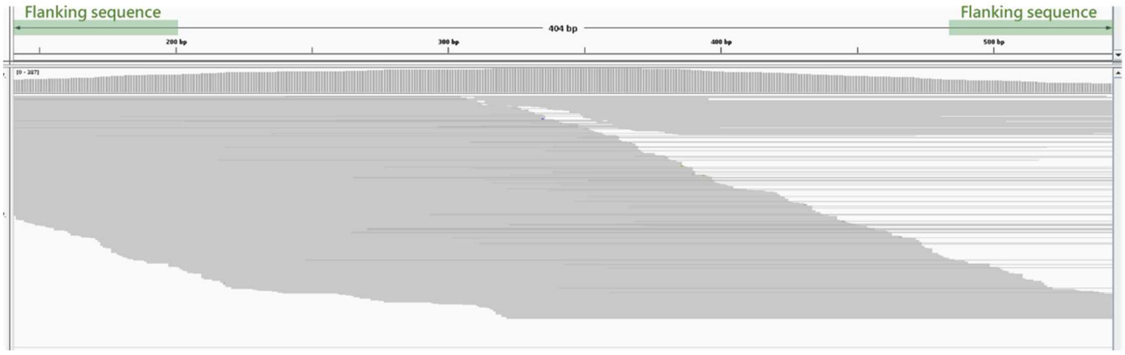

HG00673: TRAV8-4*01 & TRAV8-4*01_GWC4

TRAV8-4*01

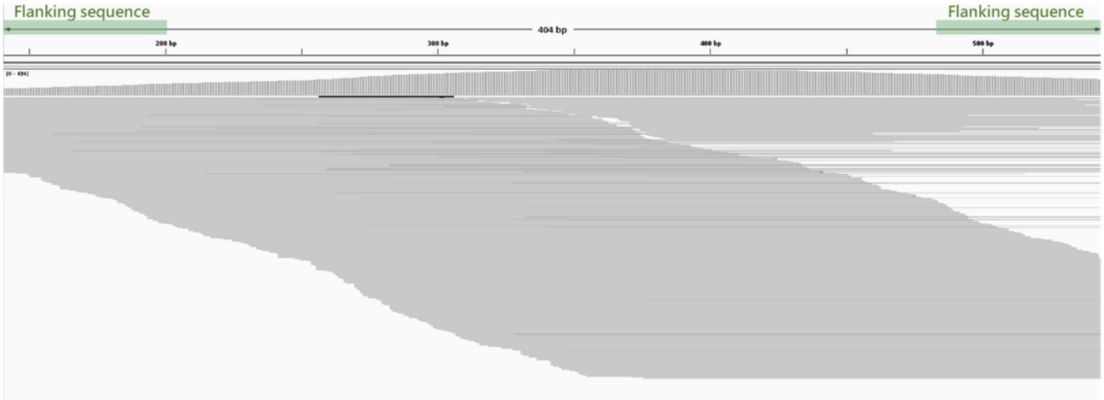


TRAV8-4*01_GWC4

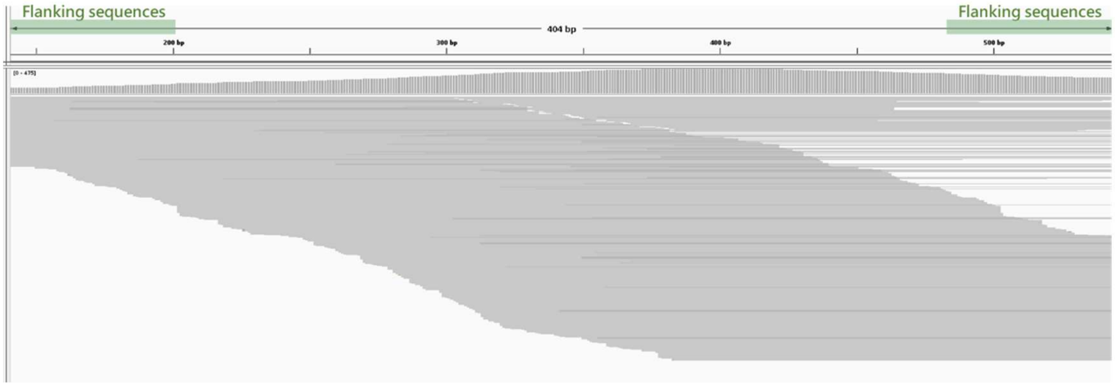


HG01891: TRAV8-4*01 & TRAV8-4*01_MEVB

TRAV8-4*01

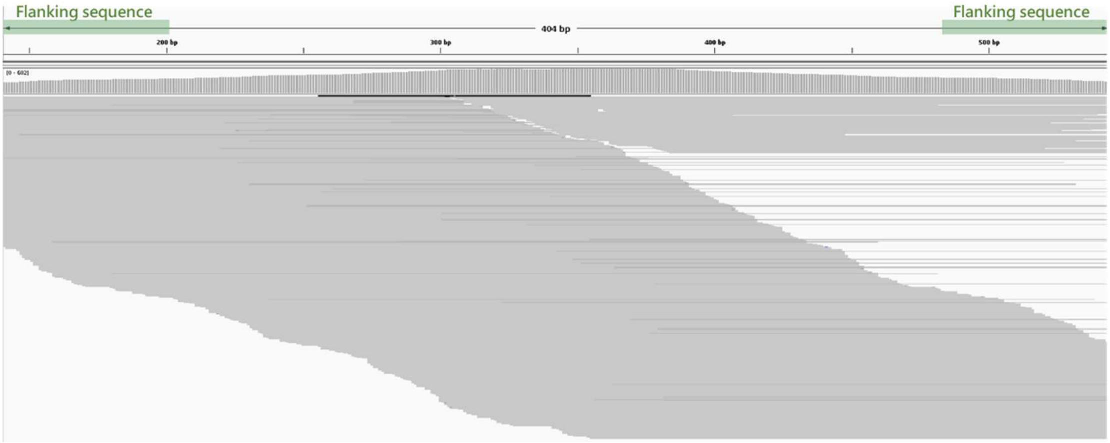

TRAV8-4*01_MEVB

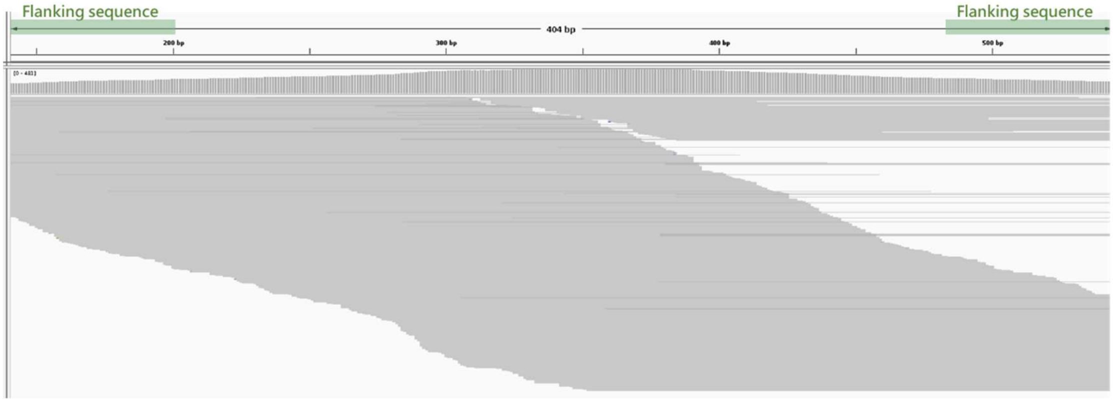


NA21309: TRAV8-4*01 & TRAV8-4*01_ODFF

TRAV8-4*01

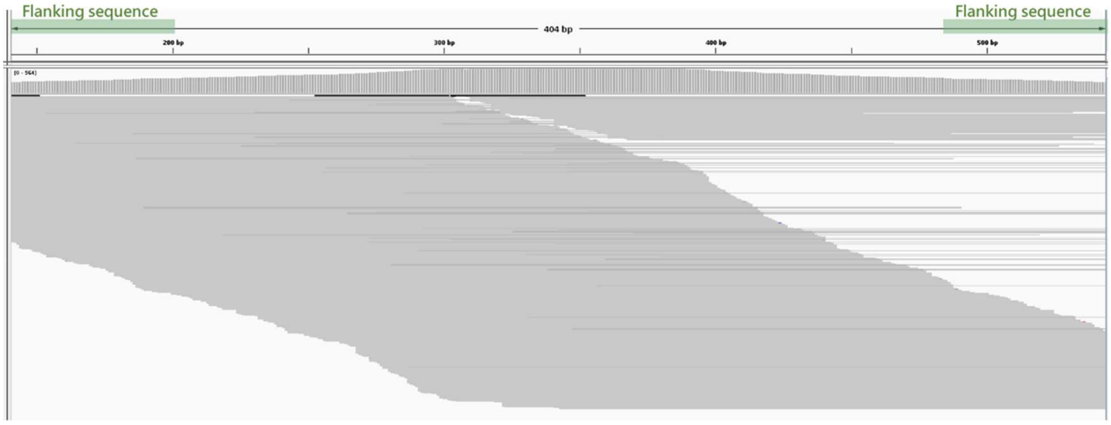


TRAV8-4*01_ODFF

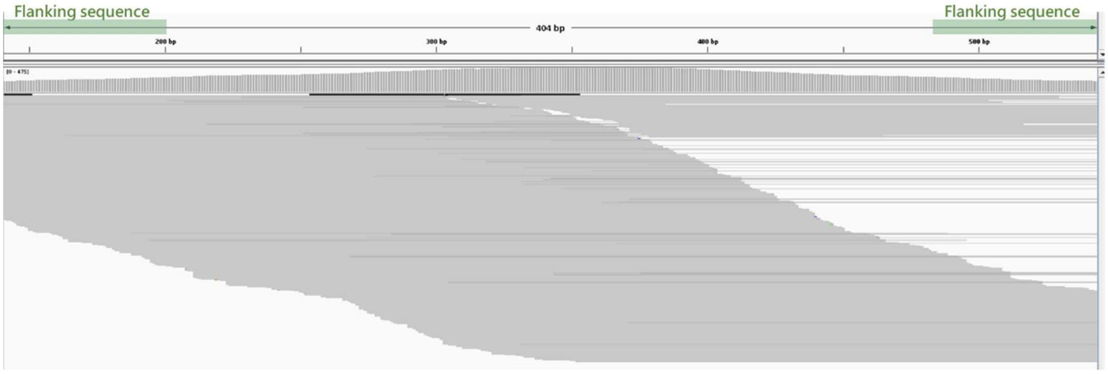


HG02717: TRAV8-4*01_NTMW & TRAV8-4*01_Q6HW

TRAV8-4*01

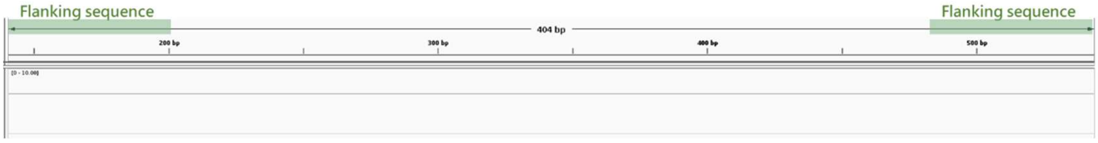

TRAV8-4*01_NTMW

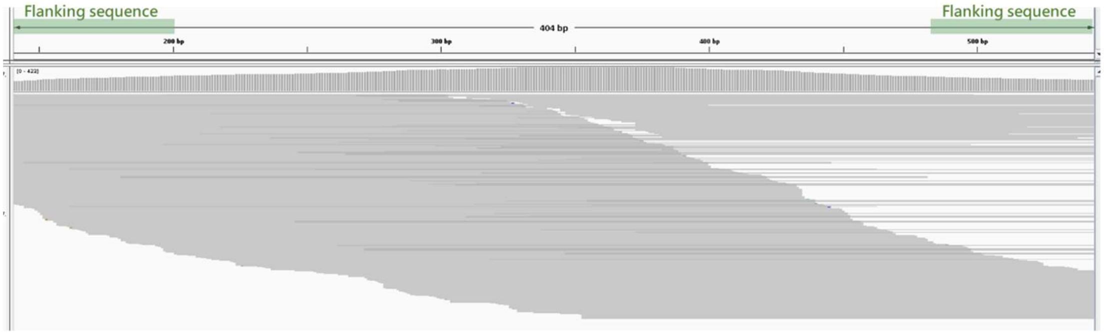


TRAV8-4*01_Q6HW

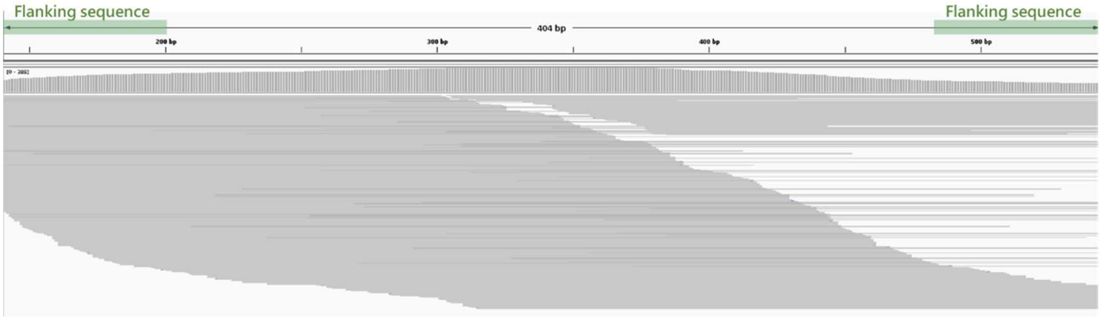


c.

TRAV14/DV4*01

```
TRAV14/DV4*01         GCCCAGAAGATAACTCAAACCCAACCAGGAATGTTCGTGCAGGAAAAGGAGGCTGTGACTCTGGACTGCACATATGACACCAGTGATCCAAGTTATGGTC    100
TRAV14/DV4*01_4XZE    ....................................................................................................
TRAV14/DV4*01_7OPH    ....................................................................................................
TRAV14/DV4*01_J3MI    ....................................................................................................
TRAV14/DV4*01_H3S5    ...........................T........................................................................

TRAV14/DV4*01         TATTCTGGTACAAGCAGCCCAGCAGTGGGGAAATGATTTTTCTTATTTATCAGGGGTCTTATGACCAGCAAAATGCAACAGAAGGTCGCTACTCATTGAA    200
TRAV14/DV4*01_4XZE    ....................................................................................................
TRAV14/DV4*01_7OPH    ....................................................................................................
TRAV14/DV4*01_J3MI    ................................................................................A...................
TRAV14/DV4*01_H3S5    ....................................................................................................

TRAV14/DV4*01         TTTCCAGAAGGCAAGAAAATCCGCCAACCTTGTCATCTCCGCTTCACAACTGGGGGACTCAGCAATGTACTTCTGTGCAATGAGAGAGGG    290
TRAV14/DV4*01_4XZE    ....................................T.....................................................
TRAV14/DV4*01_7OPH    .....................................................................T....................
TRAV14/DV4*01_J3MI    .....................................................................T....................
TRAV14/DV4*01_H3S5    .....................................................................T....................
```

HG00733: TRAV14/DV4*01 & TRAV14/DV4*01_4XZE

TRAV14/DV4*01

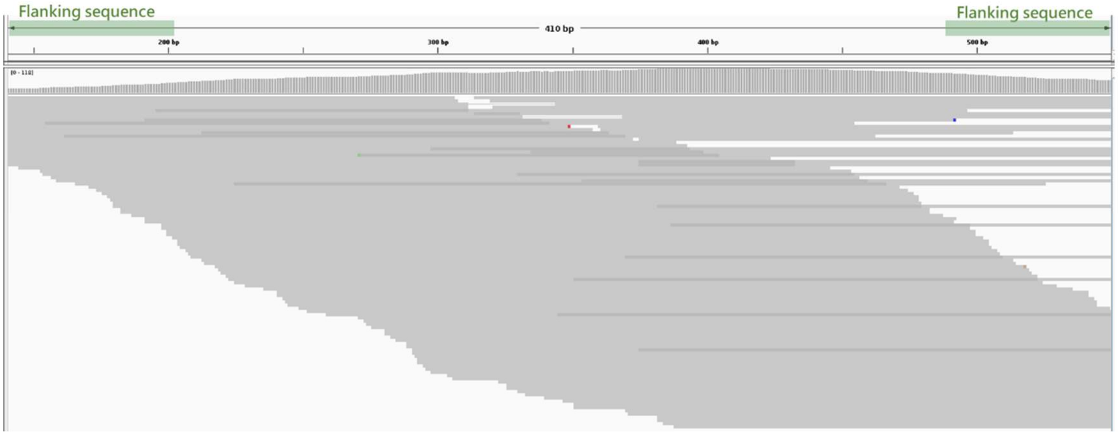

TRAV14/DV4*01_4XZE

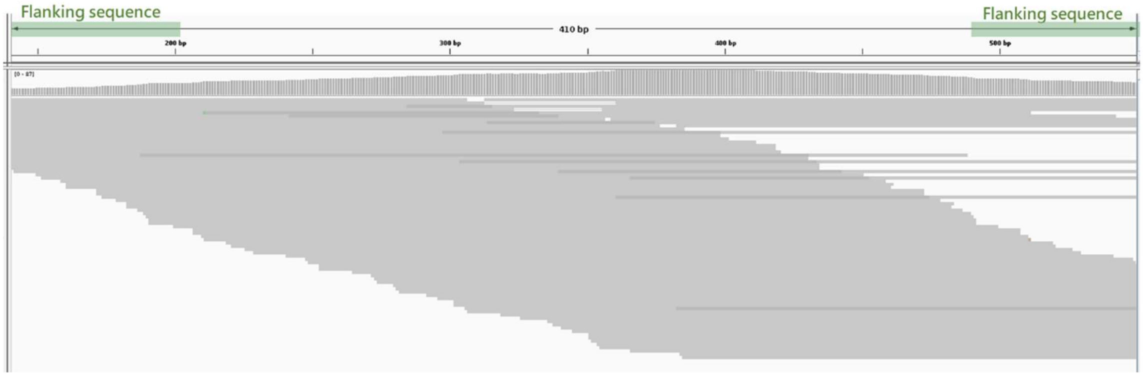


HG02717: TRAV14/DV4*01_7OPH & TRAV14/DV4*01_H3S5

TRAV14/DV4*01

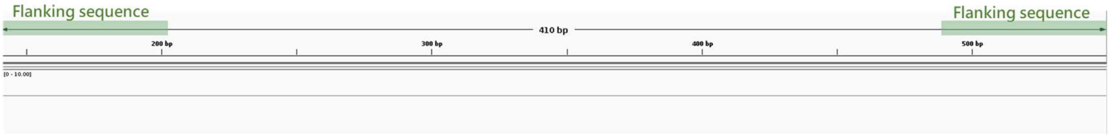


TRAV14/DV4*01_7OPH

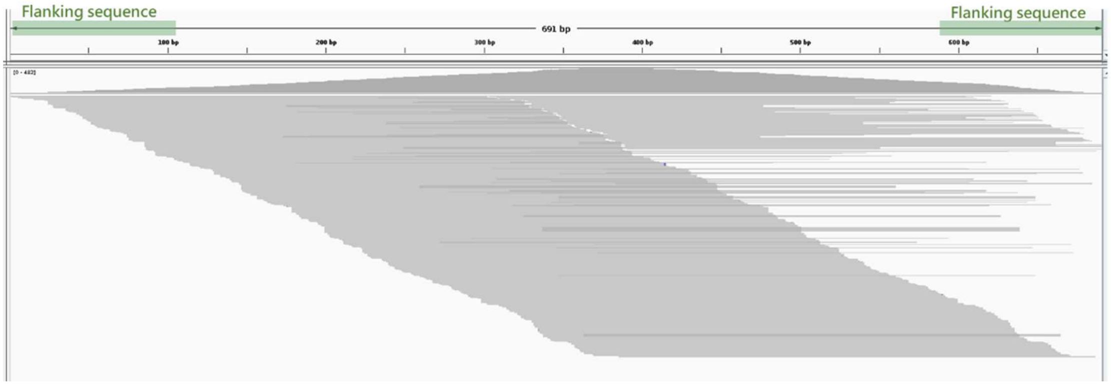


TRAV14/DV4*01_H3S5

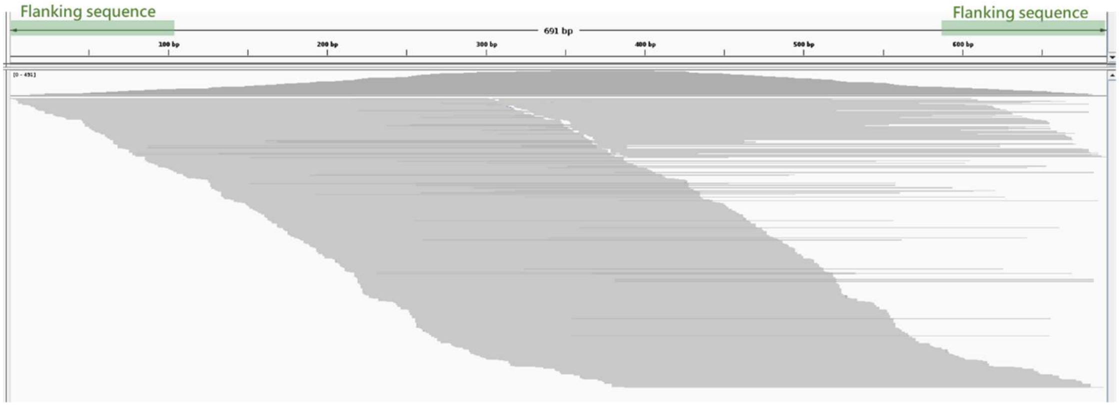

HG01891: TRAV14/DV4*01_J3MI & TRAV14/DV4*02

TRAV14/DV4*01

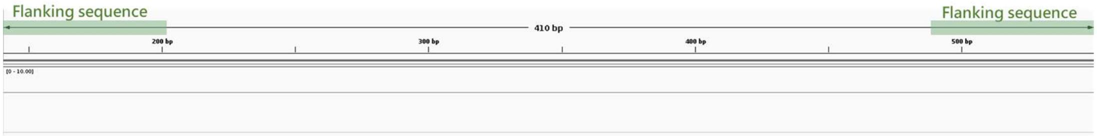


TRAV14/DV4*01_J3MI

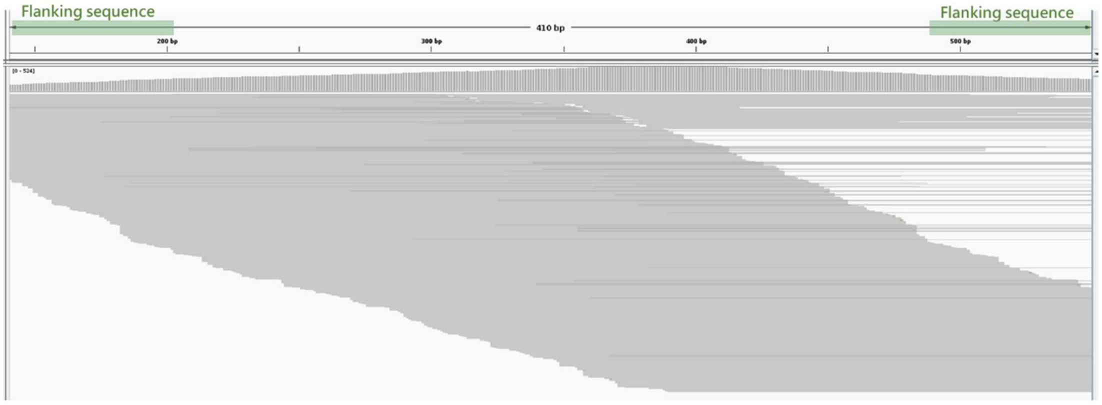


TRAV14/DV4*02

```
TRAV14/DV4*02        GCCCAGAAGATAACTCAAACCCAACCAGGAATGTTCGTGCAGGAAAAGGAGGCTGTGACTCTGGACTGCACATATGACACCAGTGATCAAAGTTATGGTC    100
TRAV14/DV4*02_COXU   ....................................................................................................
TRAV14/DV4*02_TAKJ   ...T................................................................................................

TRAV14/DV4*02        TATTCTGGTACAAGCAGCCCAGCAGTGGGGAAATGATTTTTCTTATTTATCAGGGGTCTTATGACGAGCAAAATGCAACAGAAGGTCGCTACTCATTGAA    200
TRAV14/DV4*02_COXU   .C..................................................................................................
TRAV14/DV4*02_TAKJ   ....................................................................................................

TRAV14/DV4*02        TTTCCAGAAGGCAAGAAAATCCGCCAACCTTGTCATCTCCGCTTCACAACTGGGGGACTCAGCAATGTATTTCTGTGCAATGAGAGAGGG    290
TRAV14/DV4*02_COXU   ..........................................................................................
TRAV14/DV4*02_TAKJ   ..........................................................................................
```

HG01358: TRAV14/DV4*02_COXU (homozygous)

TRAV14/DV4*02

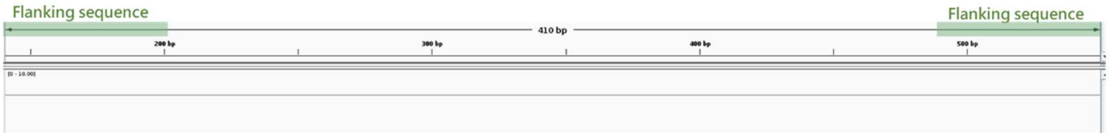


TRAV14/DV4*02_COXU

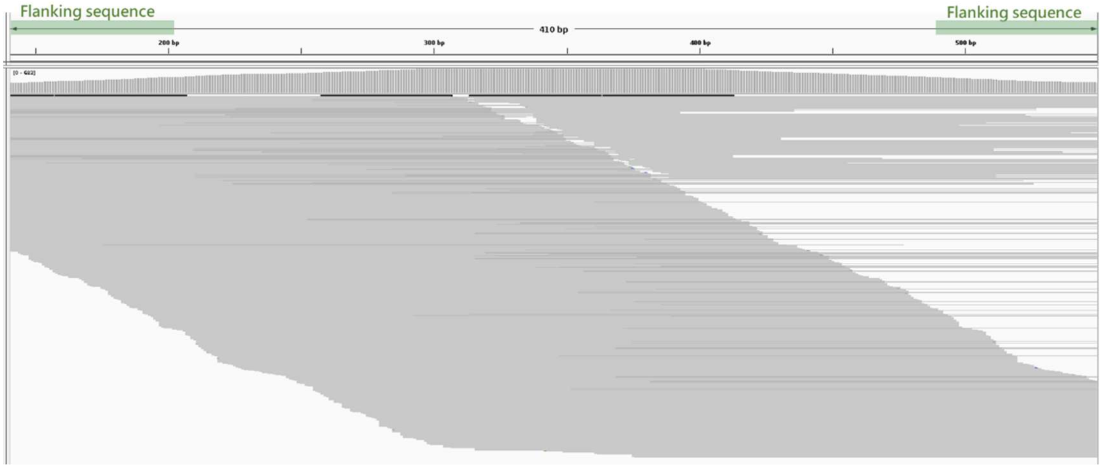

HG00673: TRAV14/DV4*01 & TRAV14/DV4*02_TAKJ

TRAV14/DV4*02

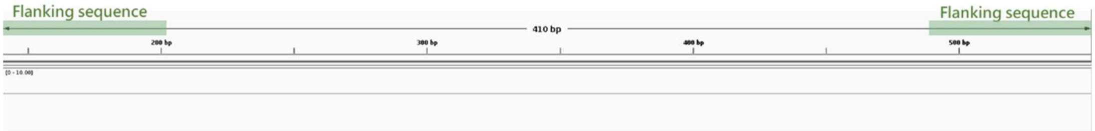


TRAV14/DV4*02_TAKJ

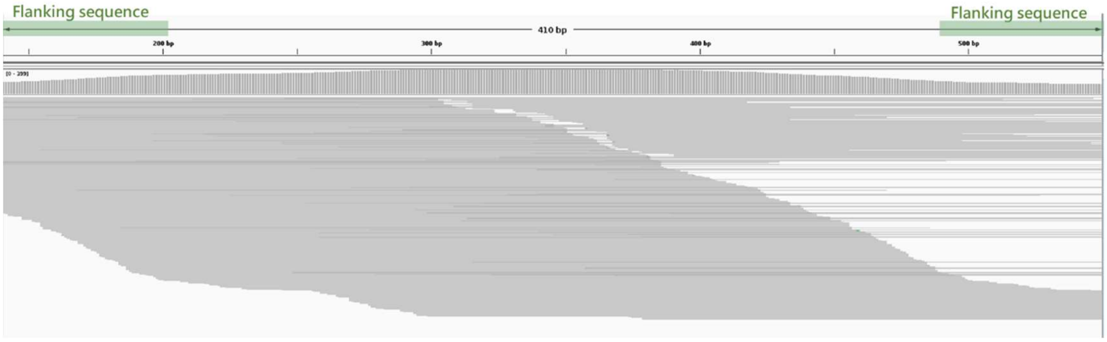


d.

TRGV3*01

```
TRGV3*01        TCTTCCAACTTGGAAGGGAGAACGAAGTCAGTCACCAGGCAGACTGGGTCATCTGCTGAAATCACTTGCGATCTTACTGTAACAAATACCTTCTACATCC   100
TRGV3*01_5AGX   ...............................................C....................................................
TRGV3*01_623Z   ....................................................................................................
TRGV3*01_XARG   ....................................................................................................
TRGV3*01_ZZNT   ....................................................................................................
TRGV3*01_GTOV   ....................................................................................................
TRGV3*01_EAGP   ........................................C...........................................................

TRGV3*01        ACTGGTACCTACACCAGGAGGGGAAGGCCCCACAGCGTCTTCTGTACTATGACGTCTCCACCGCAAGGGATGTGTTGGAATCAGGACTCAGTCCAGGAAA   200
TRGV3*01_5AGX   ....................................................................................................
TRGV3*01_623Z   ..............................................................A.....................................
TRGV3*01_XARG   ....................................................................................................
TRGV3*01_ZZNT   ....................................................................................................
TRGV3*01_GTOV   .............................................................T......................................
TRGV3*01_EAGP   ....................................................................................................

TRGV3*01        GTATTATACTCATACACCCAGGAGGTGGAGCTGGATATTGAGACTGCAAAATCTAATTGAAAATGATTCTGGGGTCTATTACTGTGCCACCTGGGACAGG   300
TRGV3*01_5AGX   ....................................................................................................
TRGV3*01_623Z   ....................................................................................................
TRGV3*01_XARG   ......C.............................................................................................
TRGV3*01_ZZNT   .....................................C..............................................................
TRGV3*01_GTOV   ....................................................................................................
TRGV3*01_EAGP   ....................................................................................................
```

HG01361: TRGV3*01 & TRGV3*01_5AGX

TRGV3*01

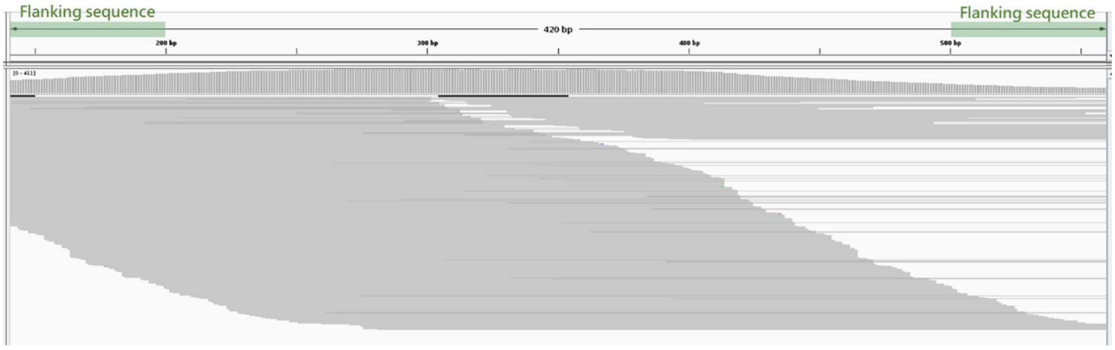

TRGV3*01_5AGX

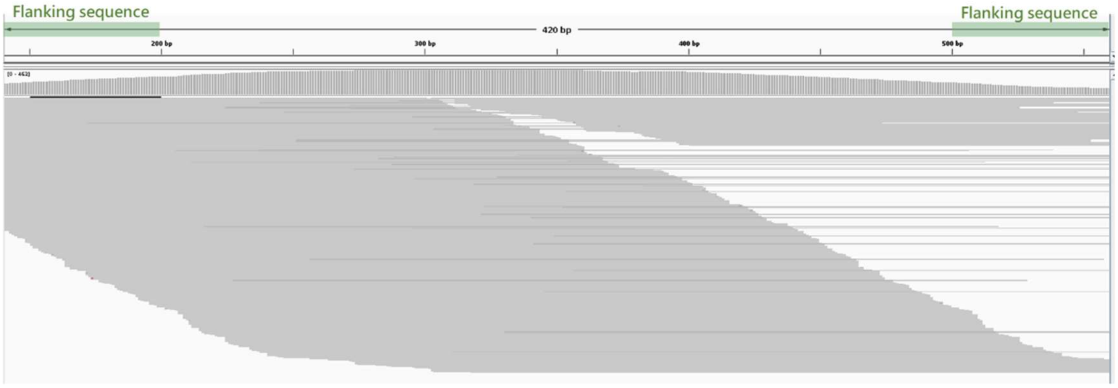


HG03516: TRGV3*01_623Z & TRGV3*01_XARG

TRGV3*01

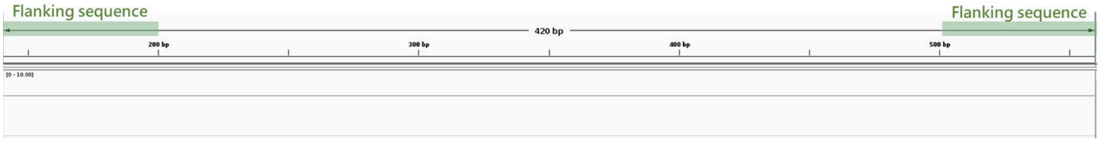


TRGV3*01_623Z

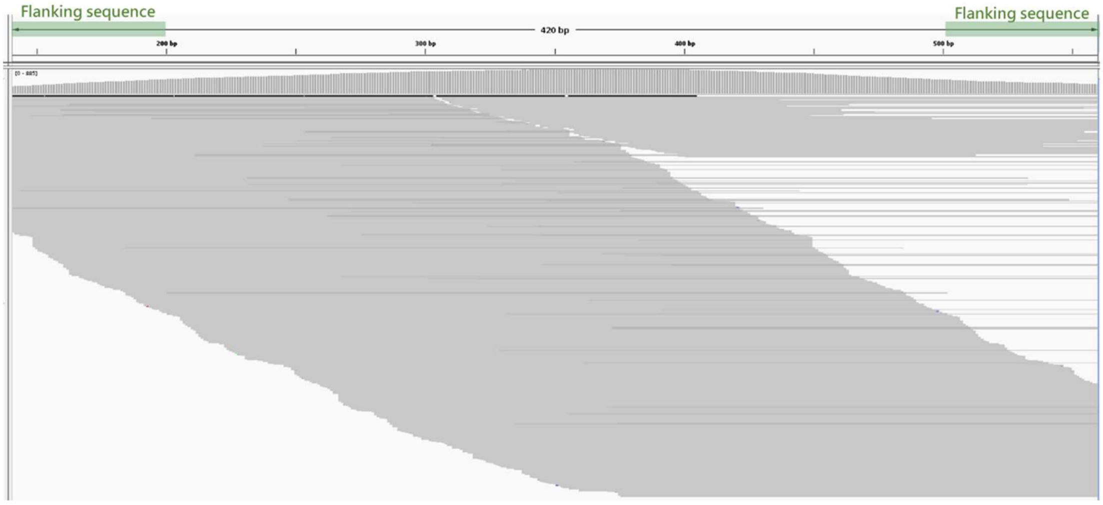


TRGV3*01_XARG

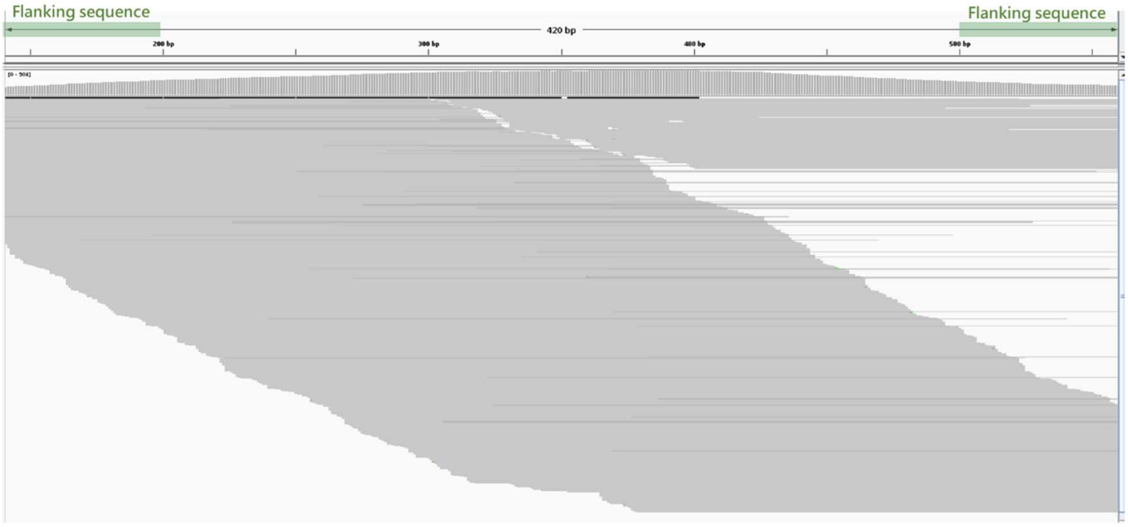

HG02559: TRGV3*01 & TRGV3*01_EAGP

TRGV3*01

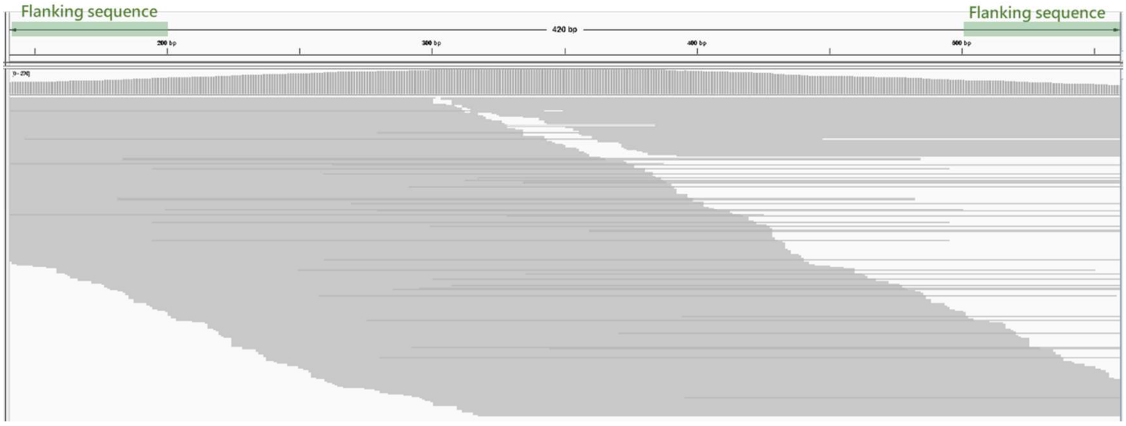


TRGV3*01_EAGP

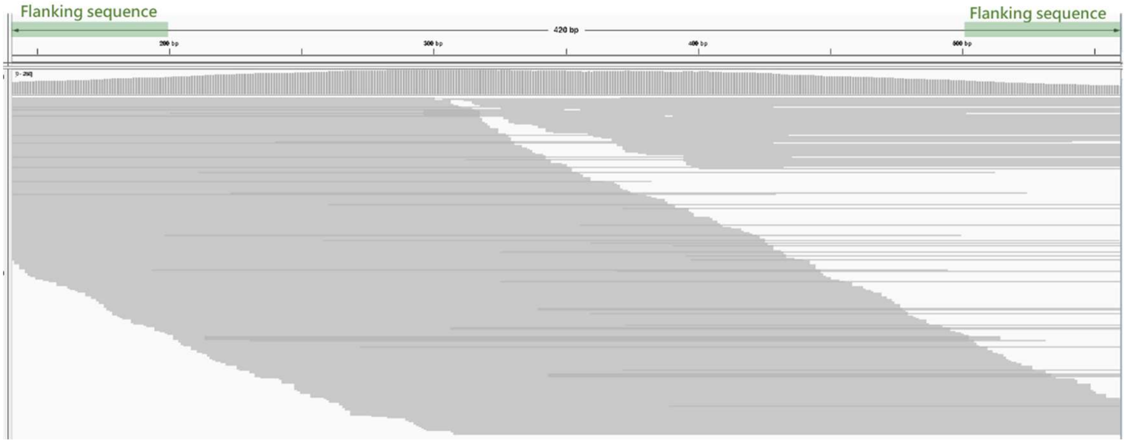


HG00673: TRGV3*01 & TRGV3*01_GTOV

TRGV3*01

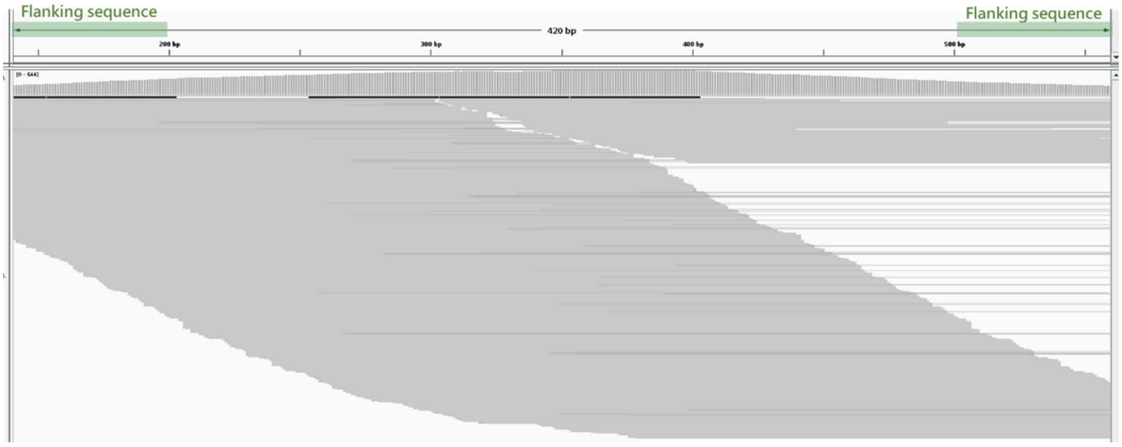

TRGV3*01_GTOV

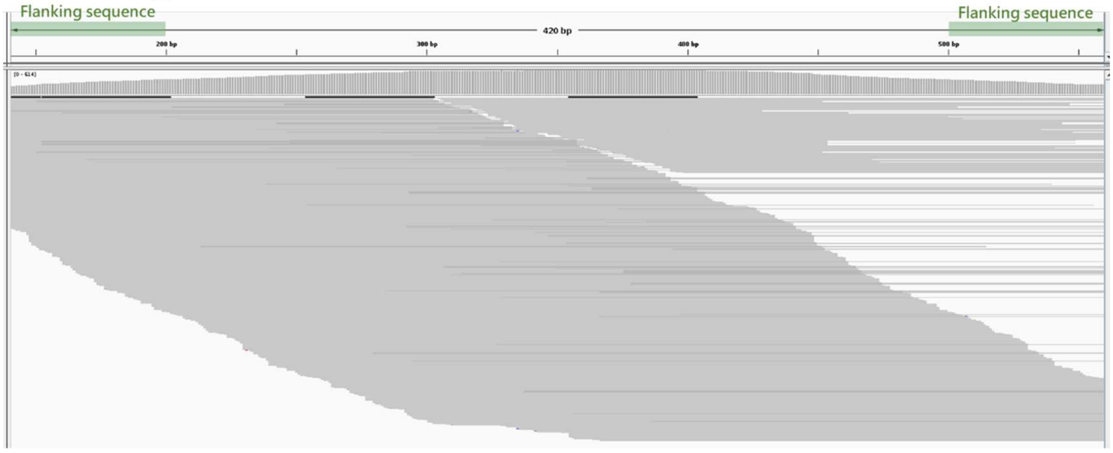


HG03540: TRGV3*01_ZZNT & TRGV3*01_XARG

TRGV3*01

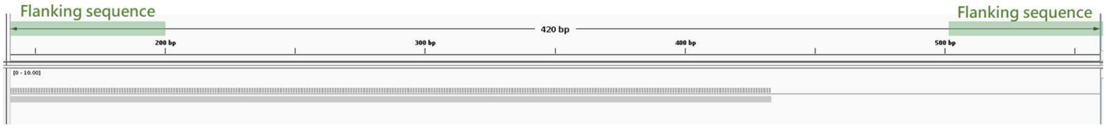


TRGV3*01_ZZNT

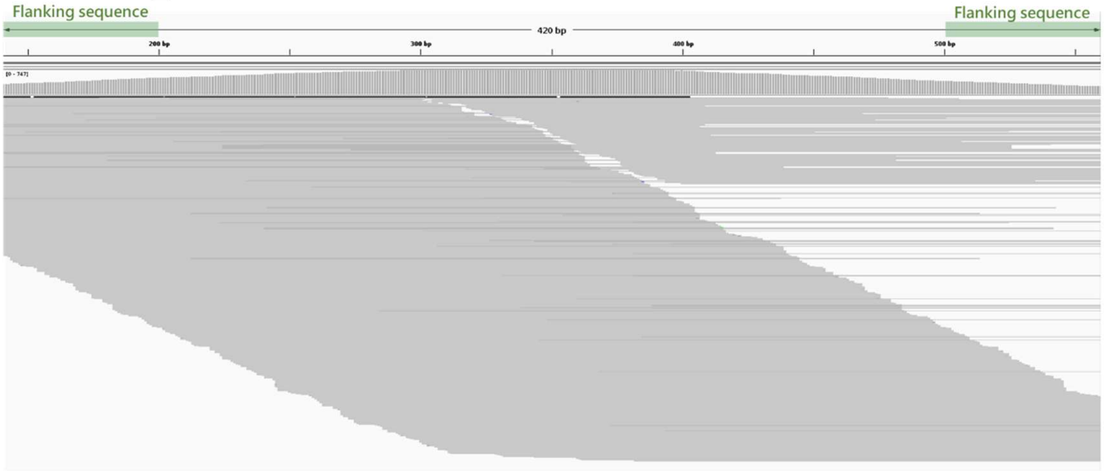


TRGV3*01_ XARG

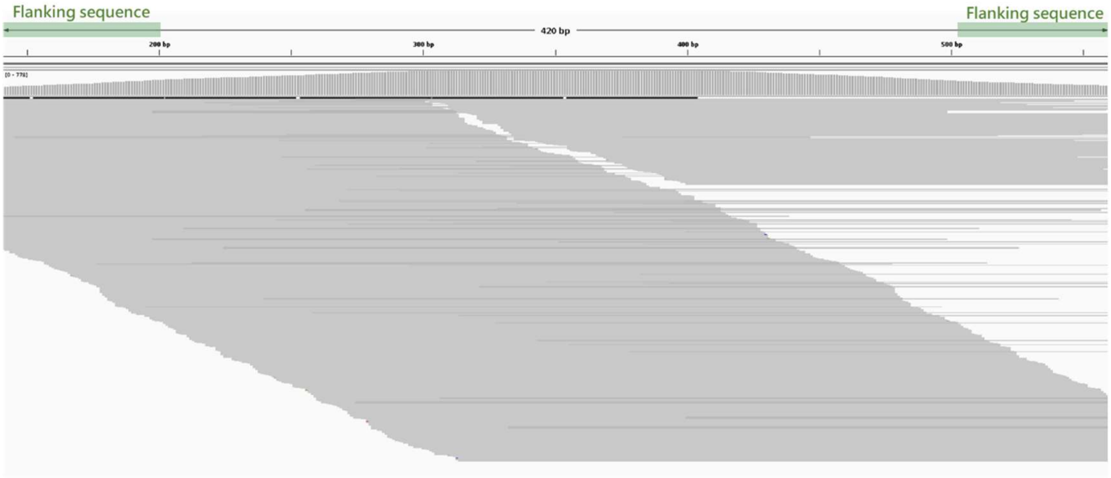

e.

### TRBV5-5*01

```
TRBV5-5*01        GACGCTGGAGTCACCCAAAGTCCCACACACCTGATCAAAACGAGAGGACAGCAAGTGACTCTGAGATGCTCTCCTATCTCTGGGCACAAGAGTGTGTCCT   100
TRBV5-5*01_A74Z   ...A................................................................................................
TRBV5-5*01_CAMA   ....................................................................................................
TRBV5-5*01_KIIP   .....................................................C..............................................
TRBV5-5*01_EABI   .....................................................C..............................................

TRBV5-5*01        GGTACCAACAGGTCCTGGGTCAGGGGCCCCAGTTTATCTTTCAGTATTATGAGAAAGAAGAGAGAGGAAGAGGAAACTTCCCTGATCGATTCTCAGCTCG   200
TRBV5-5*01_A74Z   ....................................................................................................
TRBV5-5*01_CAMA   ........G...........................................................................................
TRBV5-5*01_KIIP   ....................................................................................................
TRBV5-5*01_EABI   ....................................................................................................

TRBV5-5*01        CCAGTTCCCTAACTATAGCTCTGAGCTGAATGTGAACGCCTTGTTGCTGGGGGACTCGGCCCTGTATCTCTGTGCCAGCAGCTTGG   286
TRBV5-5*01_A74Z   ......................................................................................
TRBV5-5*01_CAMA   ......................................................................................
TRBV5-5*01_KIIP   ......................................................................................
TRBV5-5*01_EABI   ................................................A..............C......................
```

HG03098: TRBV5-5*01_A74Z & TRBV5-5*01_KIIP

TRBV5-5*01

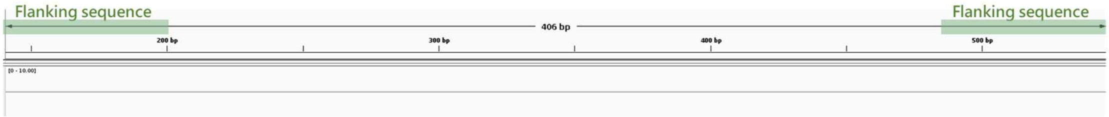


TRBV5-5*01_A74Z

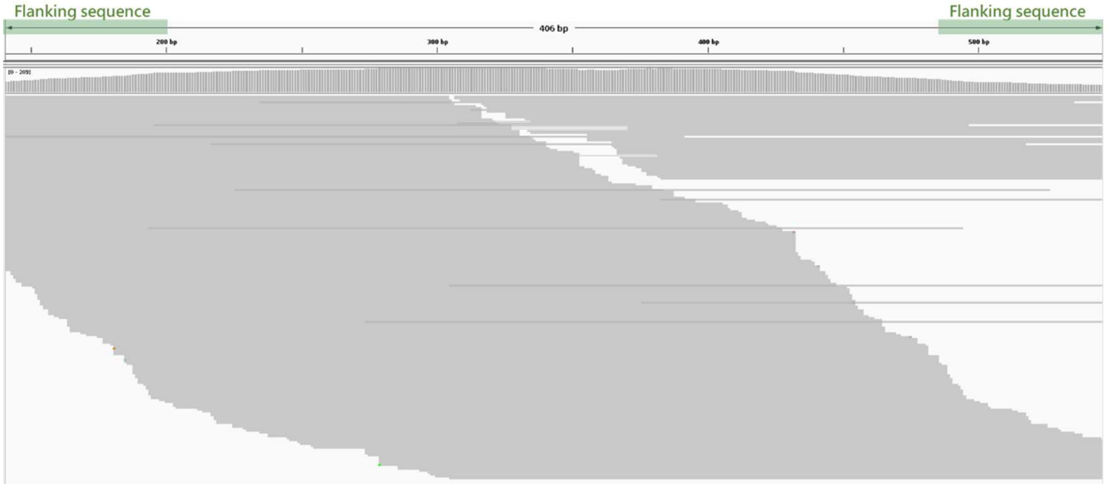


TRBV5-5*01_KIIP

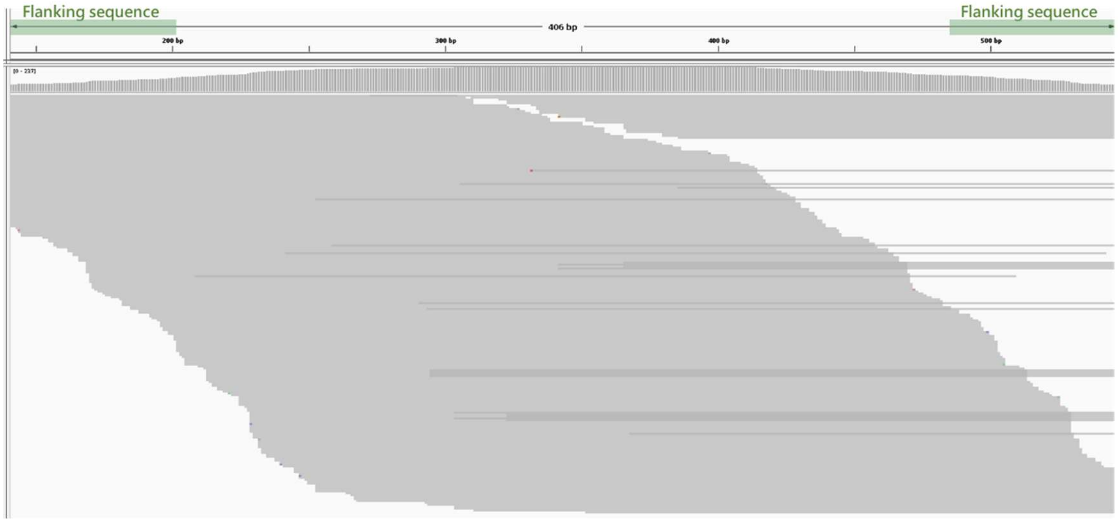

HG02055: TRBV5-5*01 & TRBV5-5*01_CAMA

TRBV5-5*01

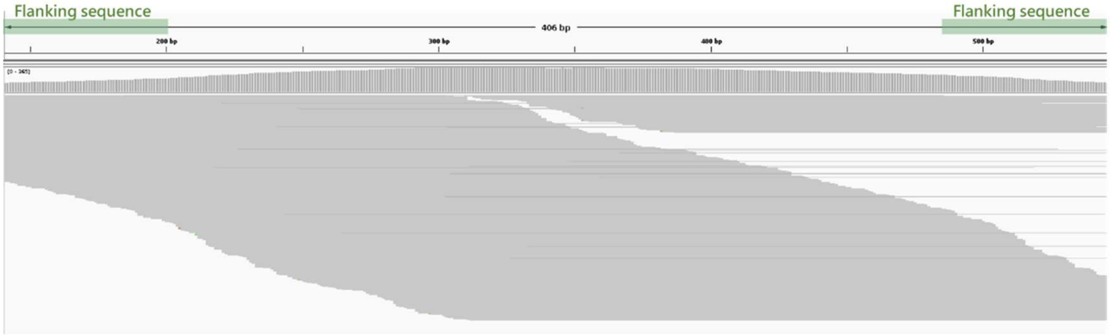


TRBV5-5*01_CAMA

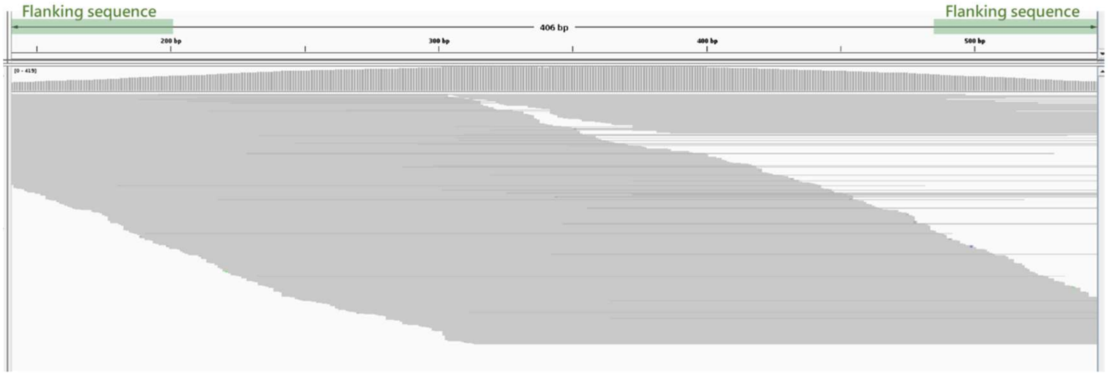


HG00673: TRBV5-5*01_EABI & TRBV5-5*01_KIIP

TRBV5-5*01

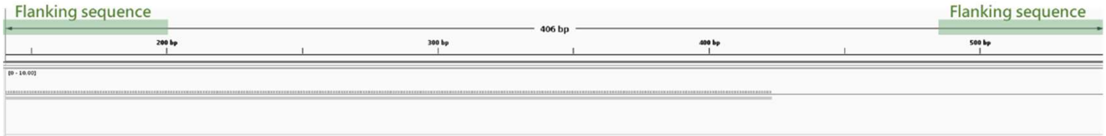


TRBV5-5*01_EABI

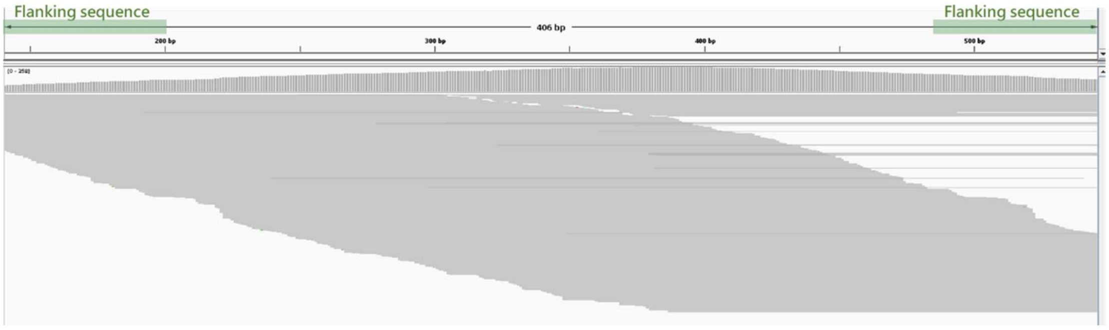

TRBV5-5*01_KIIP

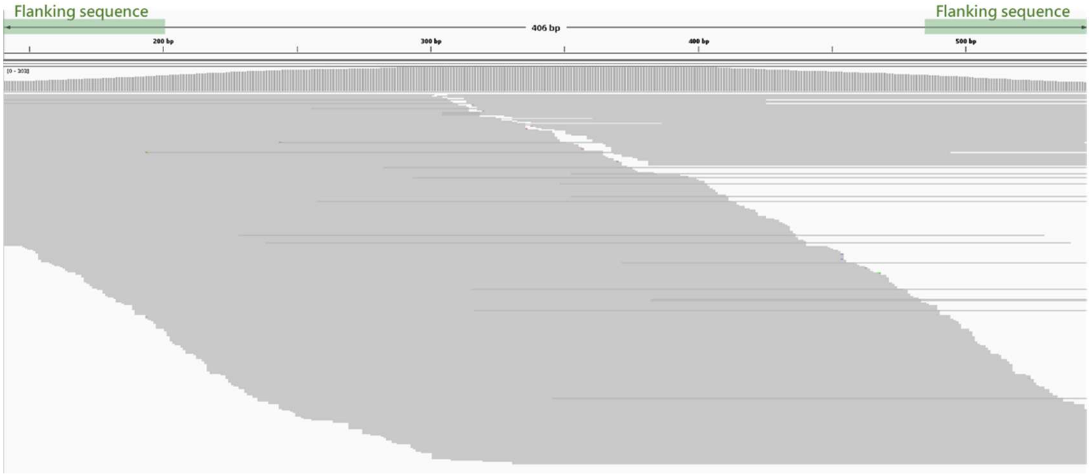


f.

TRBV5-8*01

```
TRBV5-8*01        GAGGCTGGAGTCACACAAAGTCCCACACACCTGATCAAAACGAGAGGACAGCAAGCGACTCTGAGATGCTCTCCTATCTCTGGGCACACCAGTGTGTACT   100
TRBV5-8*01_7EBE   .......................................................T............................................
TRBV5-8*01_H3GH   ....................................................................................................
TRBV5-8*01_RS7C   .......................................................T............................................
TRBV5-8*01_YC2B   .......................................................T............................................
TRBV5-8*01_AK3E   .......................................................T..........................................G.

TRBV5-8*01        GGTACCAACAGGCCCTGGGTCTGGGCCTCCAGTTCCTCCTTTGGTATGACGAGGGTGAAGAGAGAAACAGAGGAAACTTCCCTCCTAGATTTTCAGGTCG   200
TRBV5-8*01_7EBE   ....................................................................................................
TRBV5-8*01_H3GH   ....................................................................................................
TRBV5-8*01_RS7C   ....................................................................................................
TRBV5-8*01_YC2B   ..................................................................................................G.
TRBV5-8*01_AK3E   ....................................................................................................

TRBV5-8*01        CCAGTTCCCTAATTATAGCTCTGAGCTGAATGTGAACGCCTTGGAGCTGGAGGACTCGGCCCTGTATCTCTGTGCCAGCAGCTTGG   286
TRBV5-8*01_7EBE   ....................................T.................................................
TRBV5-8*01_H3GH   ....................................T.................................................
TRBV5-8*01_RS7C   ......................................................................................
TRBV5-8*01_YC2B   ....................................T.................................................
TRBV5-8*01_AK3E   ......................................................................................
```

HG00438: TRBV5-8*01_7EBE (h1 only)

TRBV5-8*01

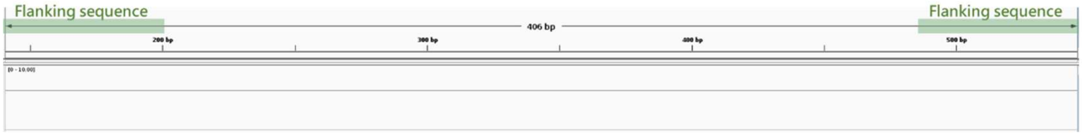


TRBV5-8*01_7EBE

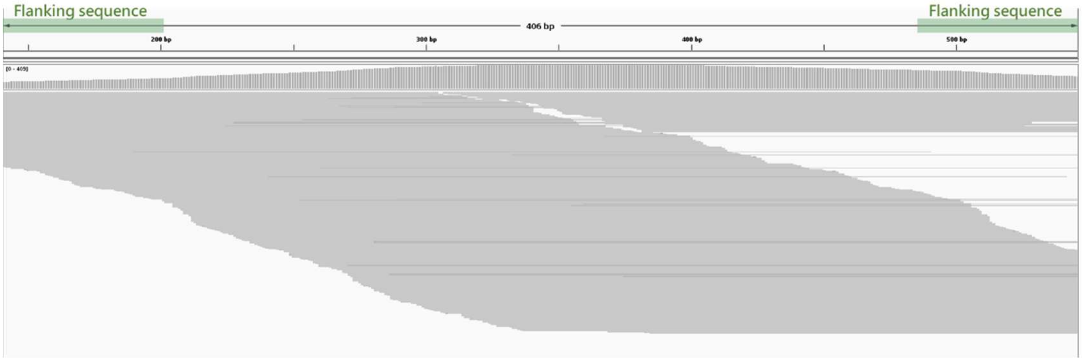

HG02055: TRBV5-8*01 & TRBV5-8*01_H3GH

TRBV5-8*01

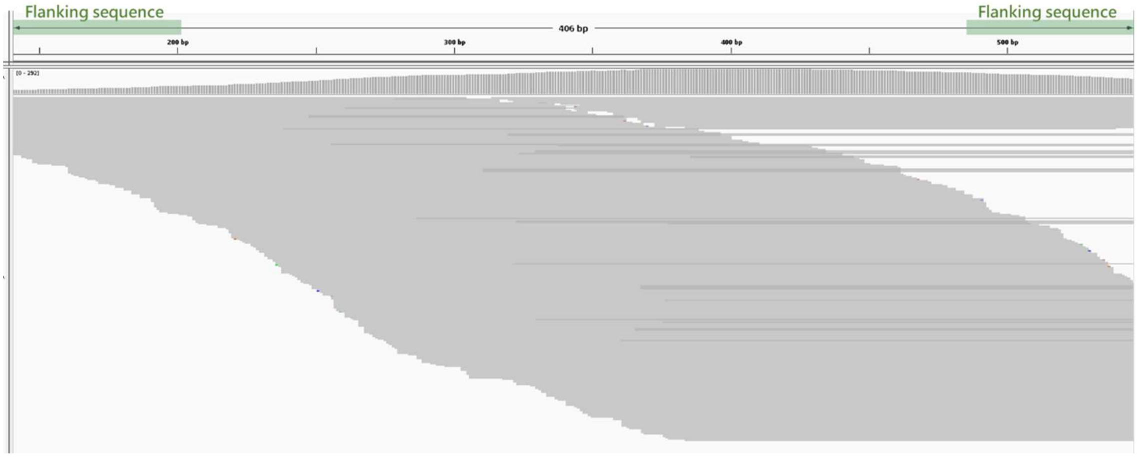


TRBV5-8*01_H3GH

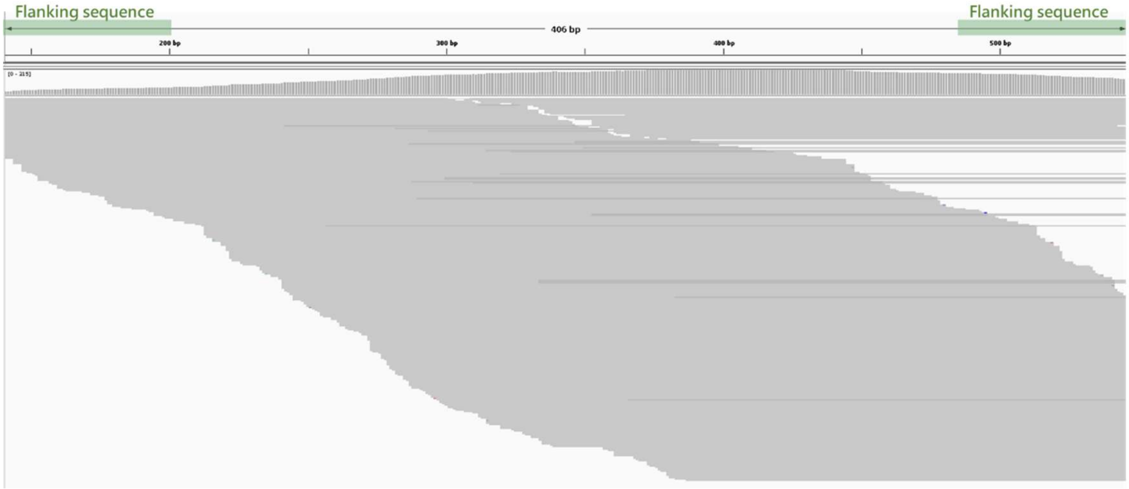


HG02486: TRBV5-8*01 & TRBV5-8*01_AK3E

TRBV5-8*01

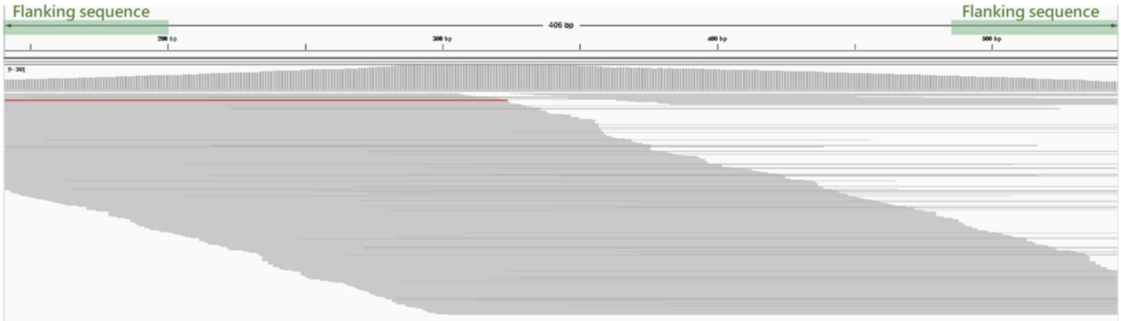

TRBV5-8*01_AK3E

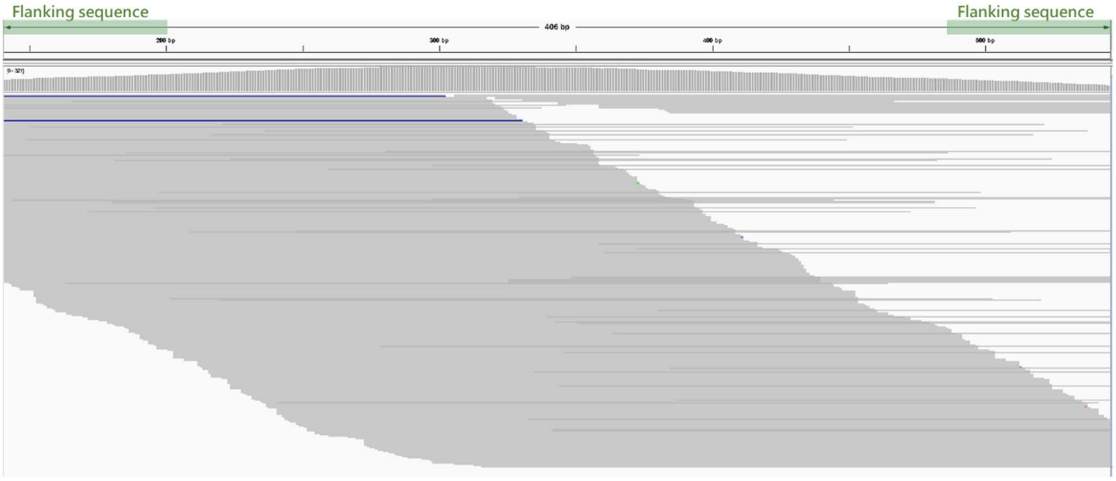


HG02257: TRBV5-8*01_RS7C (homozygous)

TRBV5-8*01

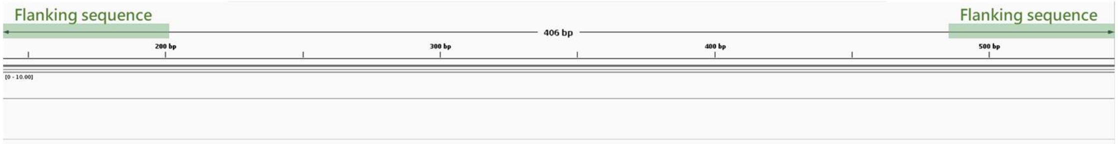


TRBV5-8*01_RS7C

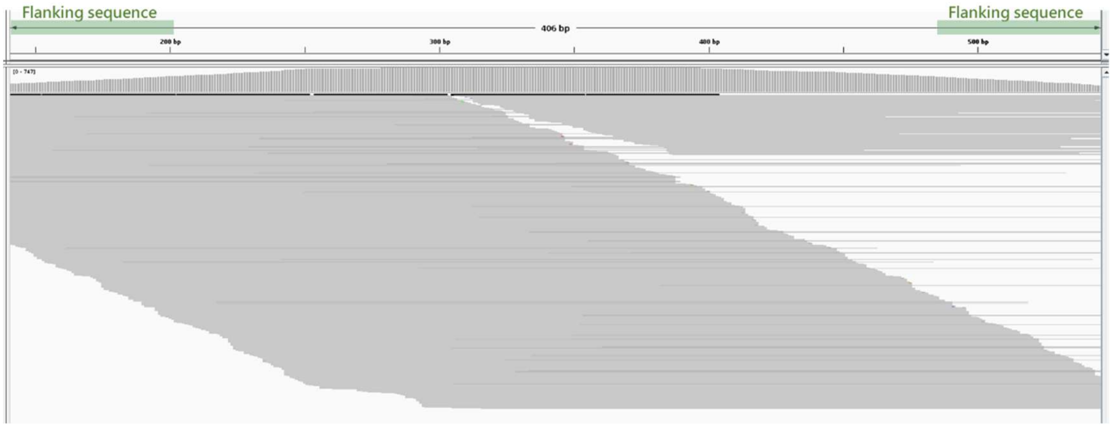

HG02717: TRBV5-8*01 & TRBV5-8*01_YC2B

TRBV5-8*01

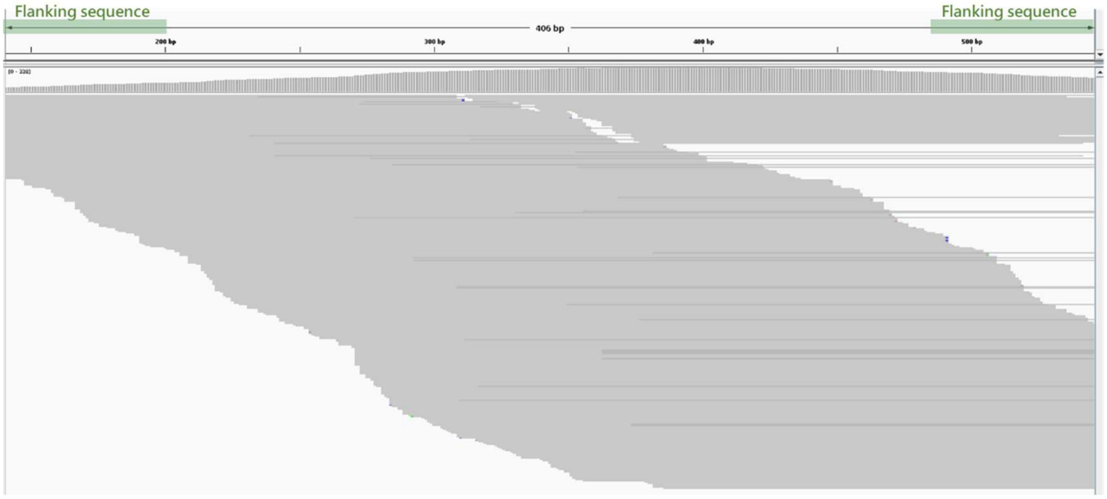


TRBV5-8*01_YC2B

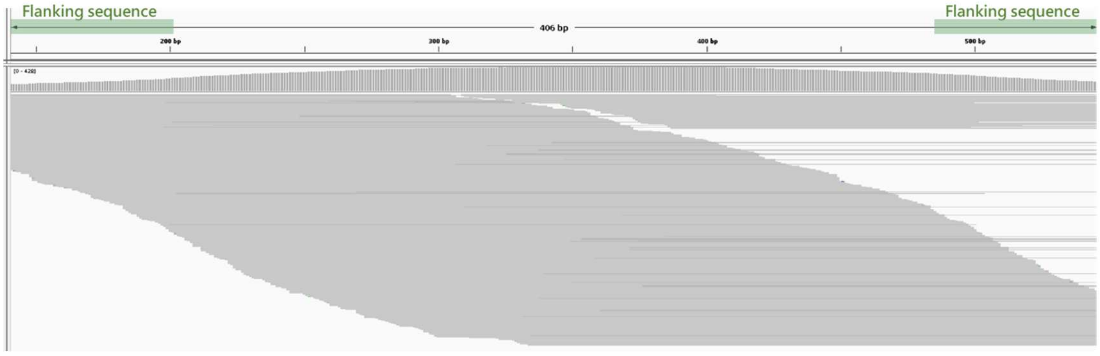


g.

TRAV12-2*01

```
TRAV12-2*01         CAGAAGGAGGTGGAGCAGAATTCTGGACCCCTCAGTGTTCCAGAGGGAGCCATTGCCTCTCTCAACTGCACTTACAGTGACCGAGGTTCCCAGTCCTTCT   100
TRAV12-2*01_7QKH    ..................................................................................A.................
TRAV12-2*01_XEQC    ....................................................................................................
TRAV12-2*01_OPDL    .....................................................................................T..............

TRAV12-2*01         TCTGGTACAGACAATATTCTGGGAAAAGCCCTGAGTTGATAATGTTCATATACTCCAATGGTGACAAAGAAGATGGAAGGTTTACAGCACAGCTCAATAA   200
TRAV12-2*01_7QKH    ....................................................................................................
TRAV12-2*01_XEQC    .............................................C......................................................
TRAV12-2*01_OPDL    .............................................C......................................................

TRAV12-2*01         AGCCAGCCAGTATGTTTCTCTGCTCATCAGAGACTCCCAGCCCAGTGATTCAGCCACCTACCTCTGTGCCGTGAACA   277
TRAV12-2*01_7QKH    .............................................................................
TRAV12-2*01_XEQC    .............................................................................
TRAV12-2*01_OPDL    .............................................................................
```

HG02257: TRAV12-2*01_7QKH & TRAV12-2*01_OPDL

TRAV12-2*01

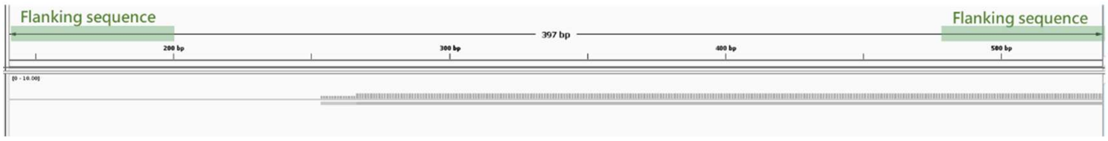


TRAV12-2*01_7QKH

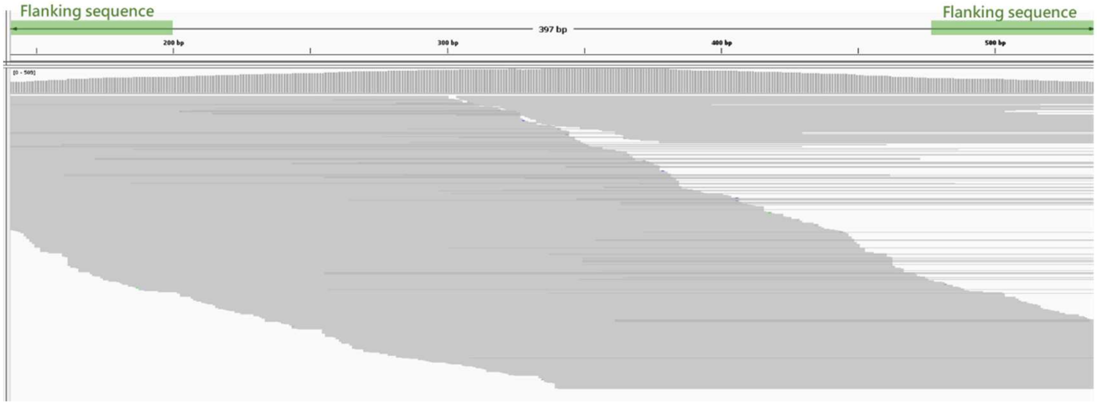


TRAV12-2*01_OPDL

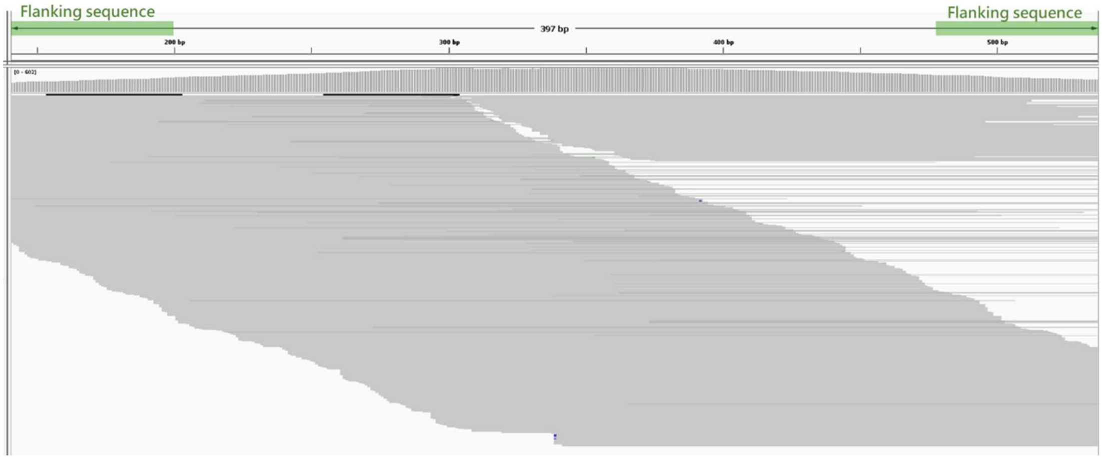


HG01928: TRAV12-2*01_XEQC (homozygous)

TRAV12-2*01

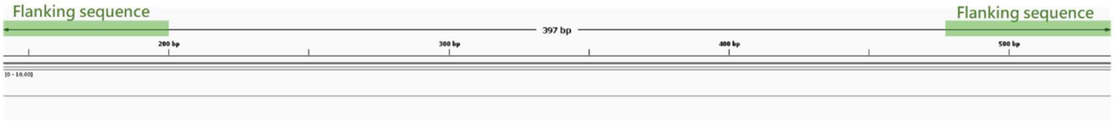

TRAV12-2*01_XEQC

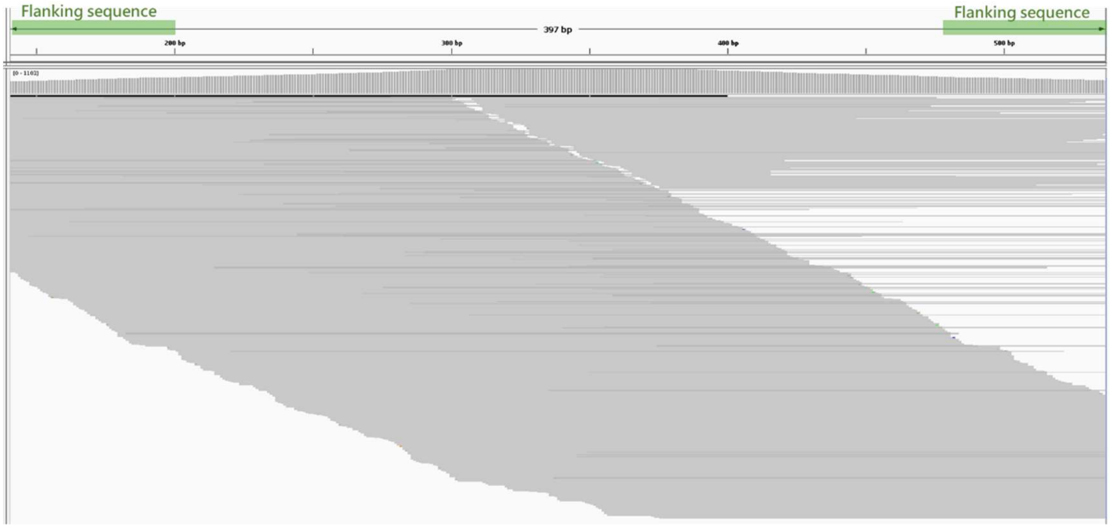


h.

TRDV3*01

```
TRDV3*01         TGTGACAAAGTAACCCAGAGTTCCCCGGACCAGACGGTGGCGAGTGGCAGTGAGGTGGTACTGCTCTGCACTTACGACACTGTATATTCAAATCCAGATT   100
TRDV3*01_HHZX    ...........................................................................A........................

TRDV3*01         TATTCTGGTACCGGATAAGGCCAGATTATTCCTTTCAGTTTGTCTTTTATGGGGATAACAGCAGATCAGAAGGTGCAGATTTTACTCAAGGACGGTTTTC   200
TRDV3*01_HHZX    ....................................................................................................

TRDV3*01         TGTGAAACACATTCTGACCCAGAAAGCCTTTCACTTGGTGATCTCTCCAGTAAGGACTGAAGACAGTGCCACTTACTACTGTGCCTTTAG   290
TRDV3*01_HHZX    ..........................................................................................
```

HG02257 TRDV3*02 & TRDV3*01_HHZX

TRDV3*01

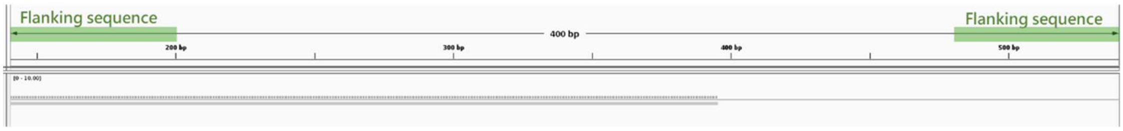


TRDV3*01_HHZX

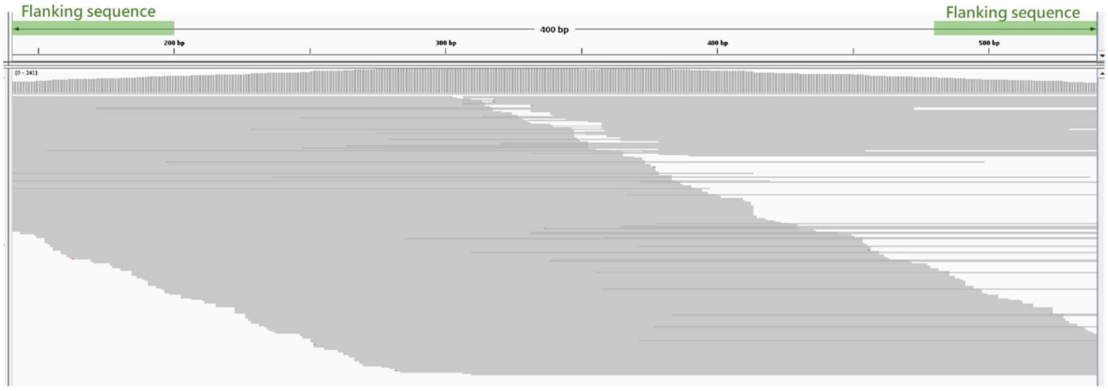

i.

TRAJ47*01

```
TRAJ47*01        TGGAATATGGAAACAAACTGGTCTTTGGCGCAGGAACCATTCTGAGAGTCAAGTCCT   57
TRAJ47*01_6LEL   ........A................................................
TRAJ47*01_ASQO   ................G........................................
TRAJ47*01_IZJW   .............................A...........................
```

HG02559: TRAJ47*01 & TRAJ47*01_6LEL

TRAJ47*01

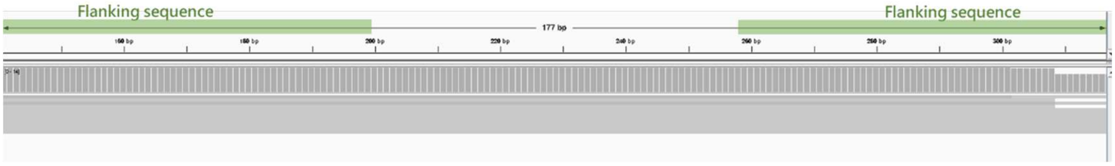


TRAJ47*01_6LEL

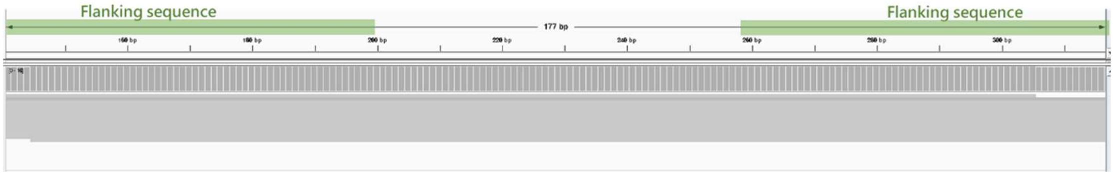


HG00673: TRAJ47*01_ASQO (homozygous)

TRAJ47*01

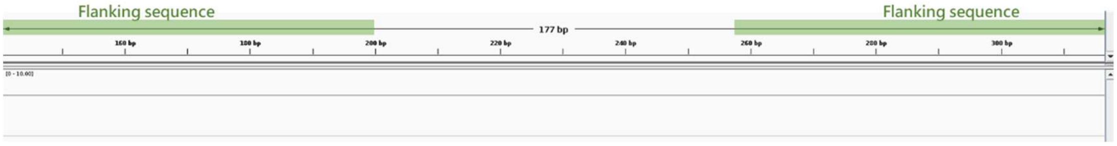


TRAJ47*01_ASQO

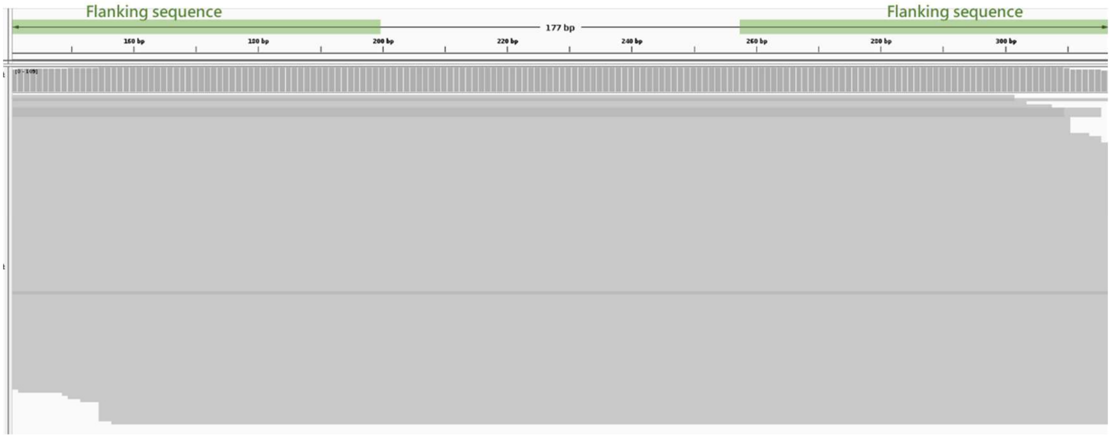

HG003516: TRAJ47*01_IZJW & TRAJ47*01_ASQO

TRAJ47*01

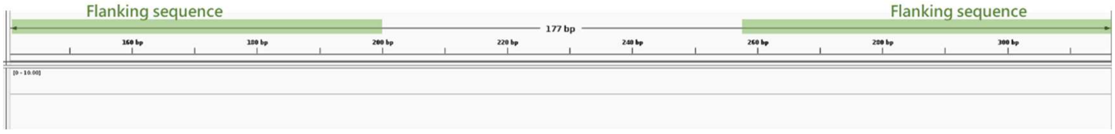


TRAJ47*01_IZJW

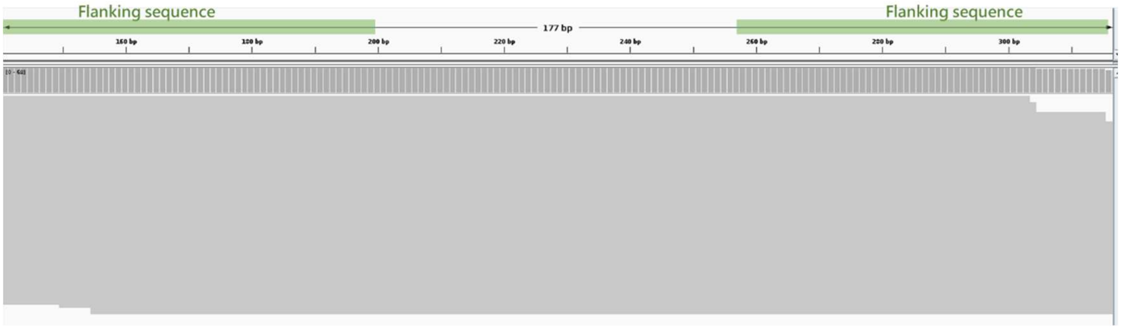


TRAJ47*01_ASQO

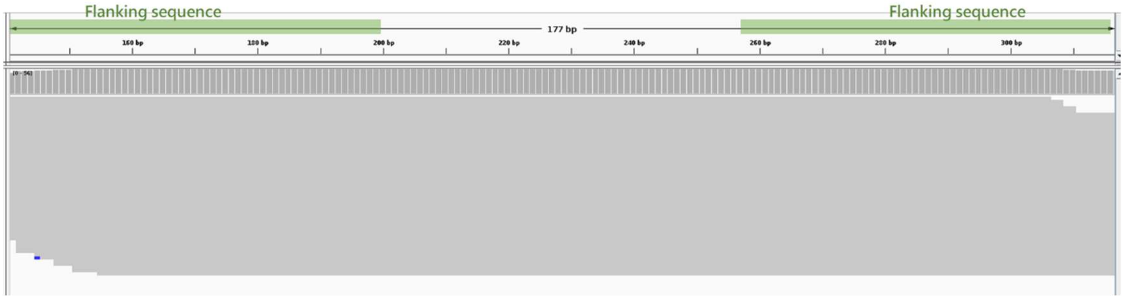


j.

TRGJP2*01

```
TRGJP2*01        ATAGTAGTGATTGGATCAAGACGTTTGCAAAAGGGACTAGGCTCATAGTAACTTCGCCTG   60
TRGJP2*01_CUJY   .....................T......................................
TRGJP2*01_NXVF   .......................................A....................
```

HG03540: TRGJP2*01 & TRGJP2*01_CUJY

TRGJP2*01

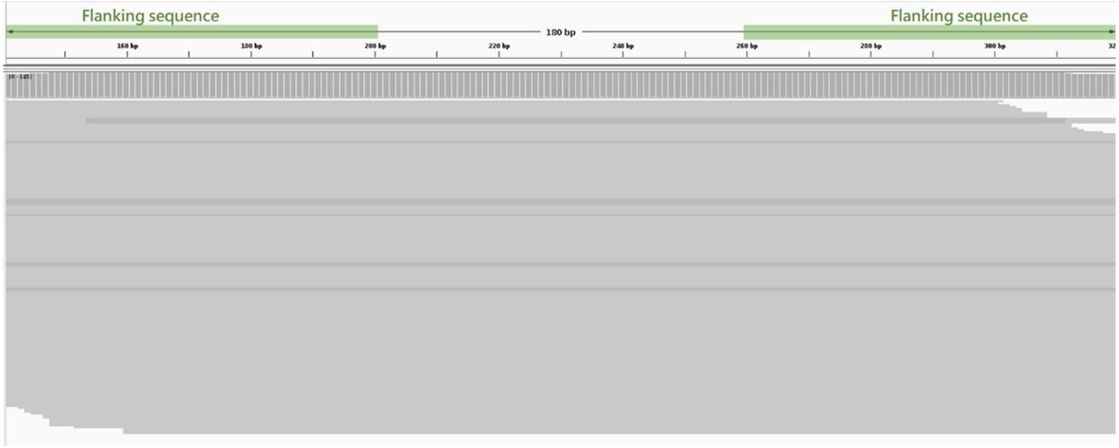

TRGJP2*01_CUJY

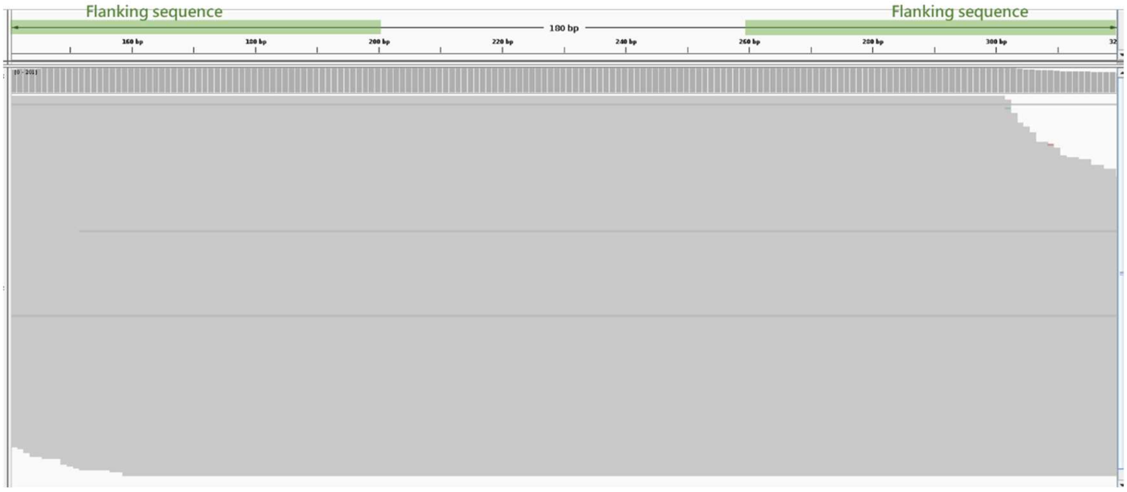


HG00673: TRGJP2*01 & TRGJP2*01_NXVF

TRGJP2*01

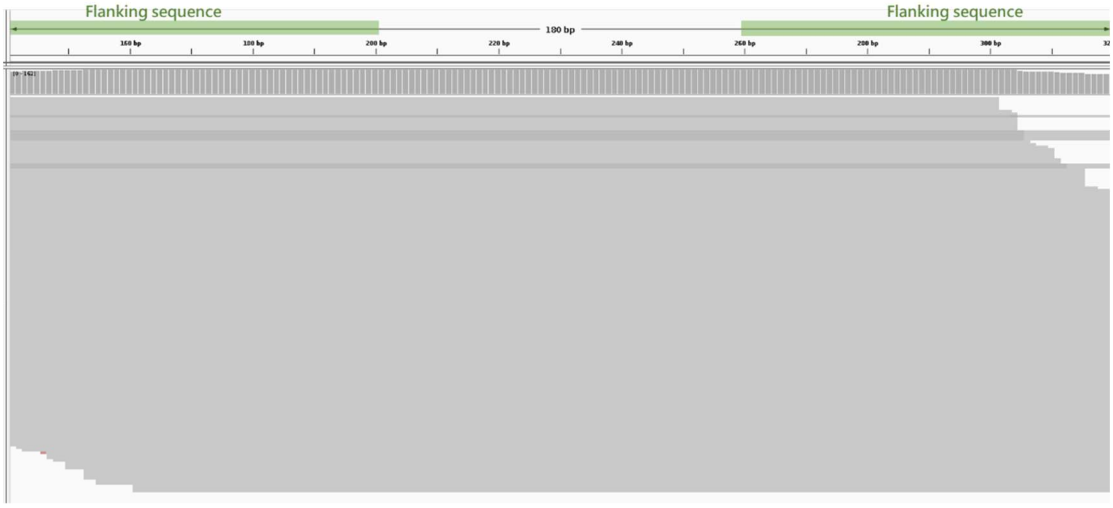


TRGJP2*01_NXVF

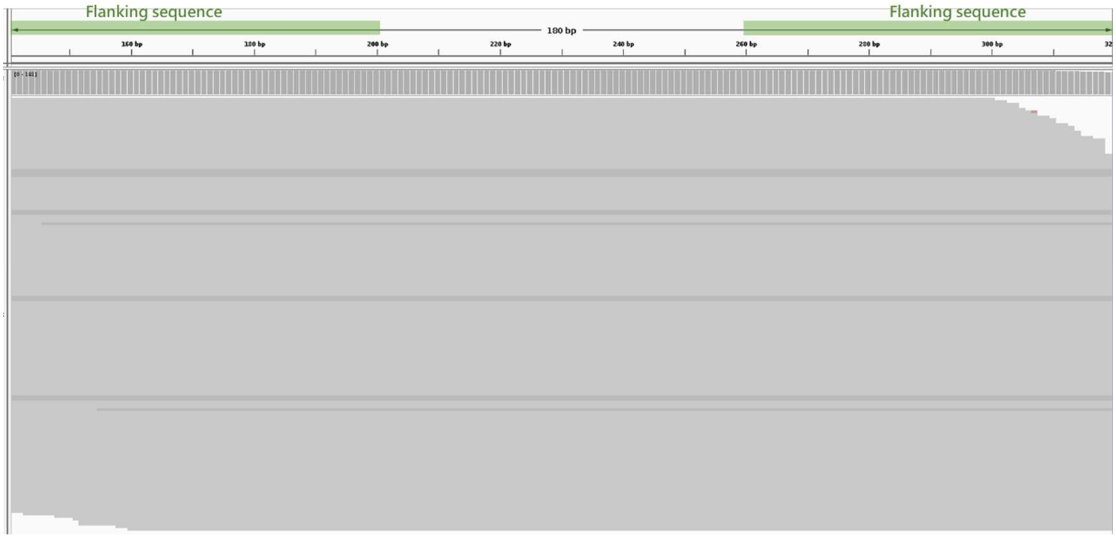

k.

TRBJ1-2*01

```
TRBJ1-2*01         CTAACTATGGCTACACCTTCGGTTCGGGGACCAGGTTAACCGTTGTAG   48
TRBJ1-2*01_WFQ7    .........................................A......
```

HG00621: TRBJ1-2*01 & TRBJ1-2*01_WFQ7

TRBJ1-2*01

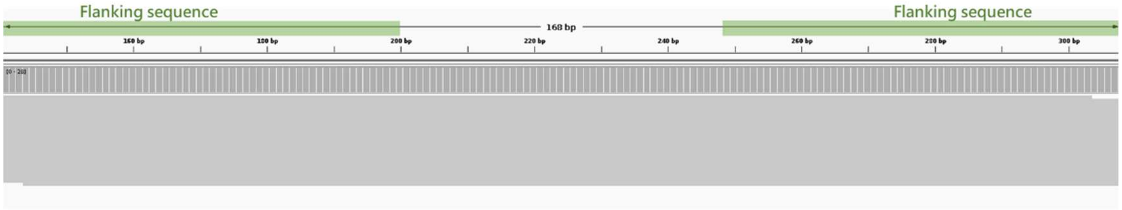


TRBJ1-2*01_WFQ7

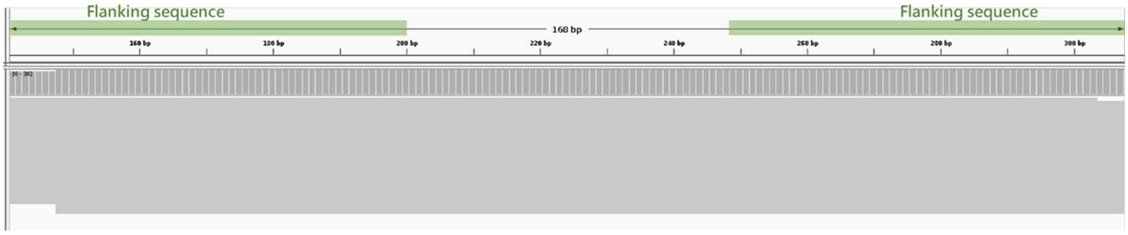


l.

TRDJ4*01

```
TRDJ4*01         CCAGACCCCTGATCTTTGGCAAAGGAACCTATCTGGAGGTACAACAAC   48
TRDJ4*01_G4F3    ...............................................T
```

HG03486: TRDJ4*01_G4F3 (homozygous)

TRDJ4*01

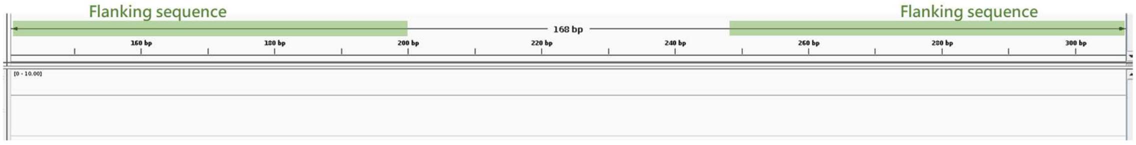

TRDJ4*01_G4F3

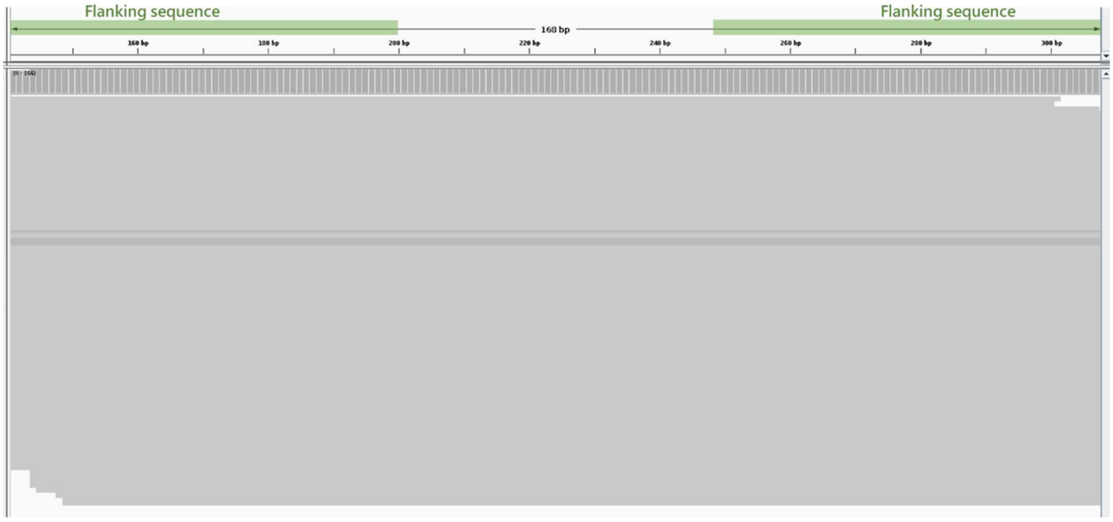


**Supplementary figure 3: Visualization of novel allele validation using IGV.**

(a) Conceptual workflow for novel allele identification and validation using the gAIRR Suite. Comparisons between alleles derived from assembly and sequencing ensure consistency and accuracy.

Validation of identified alleles is performed using sequencing data and visualized in IGV. (b - h) TRV (i - l) TRJ

a.

| Novel allele | 94 haplotypes | Frequency (%) |
|---|---|---|
| TRAV35*01_QREI | 66 | 70.21 |
| TRBV5-5*01_KIIP | 60 | 63.83 |
| TRBVC*01_O7LI | 59 | 62.77 |
| TRBV20-1*01_SXO6 | 57 | 60.64 |
| TRAV6*01_N2UE | 54 | 57.45 |
| TRBV7-5*02_3DWO | 51 | 54.26 |
| TRAV36/DV7*05_HFAO | 50 | 53.19 |
| TRAV31*01_QVZD | 48 | 51.06 |
| TRAV8-3*01_RZJS | 47 | 50 |
| TRBV24/OR9-2*03_6MOS | 47 | 50 |
| ... | ... | … |
| Total | 2466 | - |

| Novel allele | 94 haplotypes | Frequency (%) |
|---|---|---|
| TRDJ4*01_G4F3 | 33 | 35.11 |
| TRAJ47*01_ASQO | 27 | 28.72 |
| TRAJ1*01_7NLF | 10 | 10.64 |
| ... | ... | … |
| Total | 135 | - |

b.

| Allele frequency (%) | Novel allele count | |
|---|---|---|
| | TRV | TRJ |
| 50 | 10 | 0 |
| 40-49 | 6 | 0 |
| 30-39 | 10 | 1 |
| 20-29 | 18 | 1 |
| 10-19 | 39 | 1 |
| 1-9 | 222 | 27 |
| Total | 305 | 30 |

**Supplementary figure 4: Distribution and allele frequency of novel V and J alleles**
Allele frequencies were calculated from 94 haplotypes across 47 HPRC subjects.

(a) Novel V alleles (frequency >50%) and novel J alleles (frequency >10%).

(b) Distribution of novel alleles by frequency range. This summarizes the distribution of novel alleles across different frequency ranges, categorized into TRV and TRJ groups.
